# Septin Complexes Regulate Microtubule Organization and Synaptic Function at the Neuromuscular Junction

**DOI:** 10.64898/2026.03.01.708893

**Authors:** Farzaneh Larti, Taha Akkülah, Aybala Samancıoğlu, Gencay Kaan Polat, İnci Şardağ, Rüçhan Erdoğan, Arzu Çelik

## Abstract

Septins are conserved filamentous GTP-binding proteins that assemble at membranes and cytoskeletal interfaces, yet how they organize neuronal architecture *in vivo* remains incompletely understood. In neurons, microtubule organization is central to polarity, transport, and synaptic function, but the contribution of septin complexes to microtubule-dependent synaptic architecture remains unclear. Using the genetically tractable and paralog-restricted septin system of *Drosophila melanogaster*, we dissect the roles of Sep2 and Sep5 at larval neuromuscular junctions. Through integrated genetic, behavioral, immunostaining, and transcriptomic analyses, we show that septin loss disrupts pre- and postsynaptic organization and vesicle recycling while altering microtubule architecture. Notably, loss of septins shifts microtubules toward an acetylated and stabilized state, accompanied by increased expression of microtubule-associated and stabilization-linked factors, including *tau,* among the most upregulated genes and *ringmaker*, consistent with enhanced microtubule stabilization. Together, these findings position septin complexes as structural organizers that buffer microtubule state to preserve synaptic architecture, establishing septin composition as a key determinant of neuronal cytoskeletal organization *in vivo*.

## Introduction

Septins constitute a conserved family of filamentous GTP-binding proteins that associate with membrane lipids, actin filaments, and microtubules^1^. As a fourth cytoskeletal component^1^, septins assemble into nonpolar hetero-oligomeric complexes^2^ and into higher-order filaments diversified through paralog expansion^3^. This combinatorial organization generates structurally and functionally distinct assemblies, positioning septins as spatial regulators of cytoskeletal architecture^4^ rather than passive polymers. Septins selectively associate with subsets of actin filaments and microtubules to coordinate membrane organization and cytoskeletal dynamics^5–11^.

A central function of septins is to regulate microtubule networks. Distinct septin isoforms and complexes display selective affinity for microtubule lattices defined by tubulin isotype composition, nucleotide state, and post-translational modifications^8^. Septins contribute to centrosomal and Golgi-derived microtubule nucleation, suppress catastrophe, promote filament bundling, and regulate lattice stability in an assembly-dependent manner ^8^. They scaffold enzymes such as HDAC6, thereby influencing tubulin acetylation and reinforcing defined microtubule states^12^. By competing with canonical microtubule-associated proteins and localizing selectively to subsets of microtubules, septins are proposed to participate in a molecular framework that biases kinesin^13^ and dynein engagement^14^ and directs membrane trafficking^15^. In this context, septins function as determinants of microtubule stability and identity.

These functions are particularly critical in neurons, where polarity, compartmentalization, and synaptic transmission depend on highly organized microtubule arrays. Septins localize to dendritic spines, axonal growth cones, and synapses, where they influence neuronal development and function^16,17^. During neural development, septins regulate neurite outgrowth, axon guidance, and dendritic branching through coordinated interactions with actin filaments and microtubules^5,16^. In dendritic spines, SEPT7 establishes diffusion barriers at spine necks, enhancing membrane compartmentalization and synaptic stability ^17,18^. At presynaptic terminals, SEPT3 and SEPT5 regulate synaptic vesicle fusion and recycling, influencing vesicle pool organization and neurotransmitter release^19–21^. Septin misregulation impairs synaptic plasticity and disrupts axon initial segment organization, affecting neuronal polarity and excitability ^12,22,23^. Septin scaffolds likely coordinate proteins involved in vesicle trafficking and membrane dynamics at synapses^9,24^. Consistent with these roles, septin dysregulation has been linked to neurodevelopmental and neurodegenerative disorders^25^.

Despite extensive evidence that septins interface with membranes and cytoskeletal networks, the *in vivo* consequences of septin loss within intact neuronal circuits remain incompletely defined. It remains unclear whether observed phenotypes primarily reflect disrupted membrane scaffolding or altered actin and microtubule dynamics. Many mechanistic insights derive from *in vitro* systems that do not fully recapitulate neuronal architecture and synaptic connectivity. Interpretation is further complicated by heteropolymeric assembly, paralog redundancy, and post-translational regulation, which obscure subunit-specific contributions^1,26^. *In vivo* analyses are therefore required to define septin-dependent regulation of neuronal cytoskeletal organization.

The *Drosophila* genome encodes five septins, Pnut, Sep1, Sep2, Sep4, and Sep5, distributed across three subgroups^27,28^. These proteins assemble into hetero-oligomeric complexes that support membrane organization and cytoskeletal structure *in vivo*. Sep2 and its retrogene Sep5 (human SEPT6-subgroup) exhibit partial redundancy^29^ but remain insufficiently characterized in neuronal cytoskeletal regulation. *Drosophila* septins localize to sites of membrane remodeling, such as cleavage furrows^30^, and coordinate actin–microtubule cohesion during wound closure^31^ and collective cell migration ^32–34^. Structural and biochemical studies indicate that NC/G-domain interactions and subgroup interchangeability define filament architecture and functional output^35,36^.

Here, using an in vivo model with only five septin genes, we investigate the contributions of Sep2 and Sep5 to neuronal cytoskeletal organization at the *Drosophila* larval neuromuscular junction. Using genetic analysis, allelic variants, molecular modeling, behavioral assays, quantitative immunostaining, and transcriptomic profiling, we assess how septin loss affects synaptic organization. Septin disruption alters synaptic structural architecture and vesicle recycling and is associated with increased acetylated tubulin and enhanced microtubule stabilization. These cytoskeletal changes correlate with altered locomotor performance and with transcriptional modulation of microtubule and actin-binding proteins. Together, these findings identify septins as a determinant of neuronal microtubule state and synaptic structural organization *in vivo*.

## Results

### Sep2 and Sep5 are broadly expressed in the nervous system and colocalize with core septin components and microtubules

To investigate the distribution of Sep2 in the *Drosophila* nervous system, we utilized a previously established Sep2::EGFP transgenic line^37^. Confocal microscopy of third-instar larval brains and ventral nerve cords (VNC) demonstrated that Sep2::EGFP was extensively expressed throughout the nervous tissue, including neuronal somata, axonal tracts (Supp Fig. 1A-C), and synaptic terminals (Fig 1A). Sep2::EGFP exhibited enrichment in the brain lobes and along axonal projections within the VNC, where it colocalized with α-tubulin (Supp Fig. 1A). Within the central nervous system, Sep2::EGFP displayed a punctate pattern (Supp Fig. 1B). Due to the lack of Sep2-specific antibodies, its localization was confirmed through co-staining with anti-Pnut, an antibody recognizing the core dimer of *Drosophila* septin hexamer without any replaceable paralog. The overlap of Sep2::EGFP signals with Pnut in neuronal somata supports its incorporation into septin filaments (Supp Fig. 1B). Considering the known interactions between septins and microtubules, we further examined their colocalization with α-tubulin, observing Sep2::EGFP in axons, including motor neuron projections (Supp Fig. 1C). Additionally, to assess whether Sep5 exhibits similar localization, we expressed GFP-tagged Sep5 in neurons under the control of the pan-neuronal driver nSyb-Gal4. Sep5::GFP displayed a comparable distribution, with presence in cell bodies and axons, overlapping substantially with α-tubulin (Supp Fig. 1D). Collectively, these findings indicate that Sep2 is broadly expressed in the *Drosophila* nervous system and closely associates with core septin components and microtubules, supporting their involvement in neuronal structure and function.

**Figure 1.**
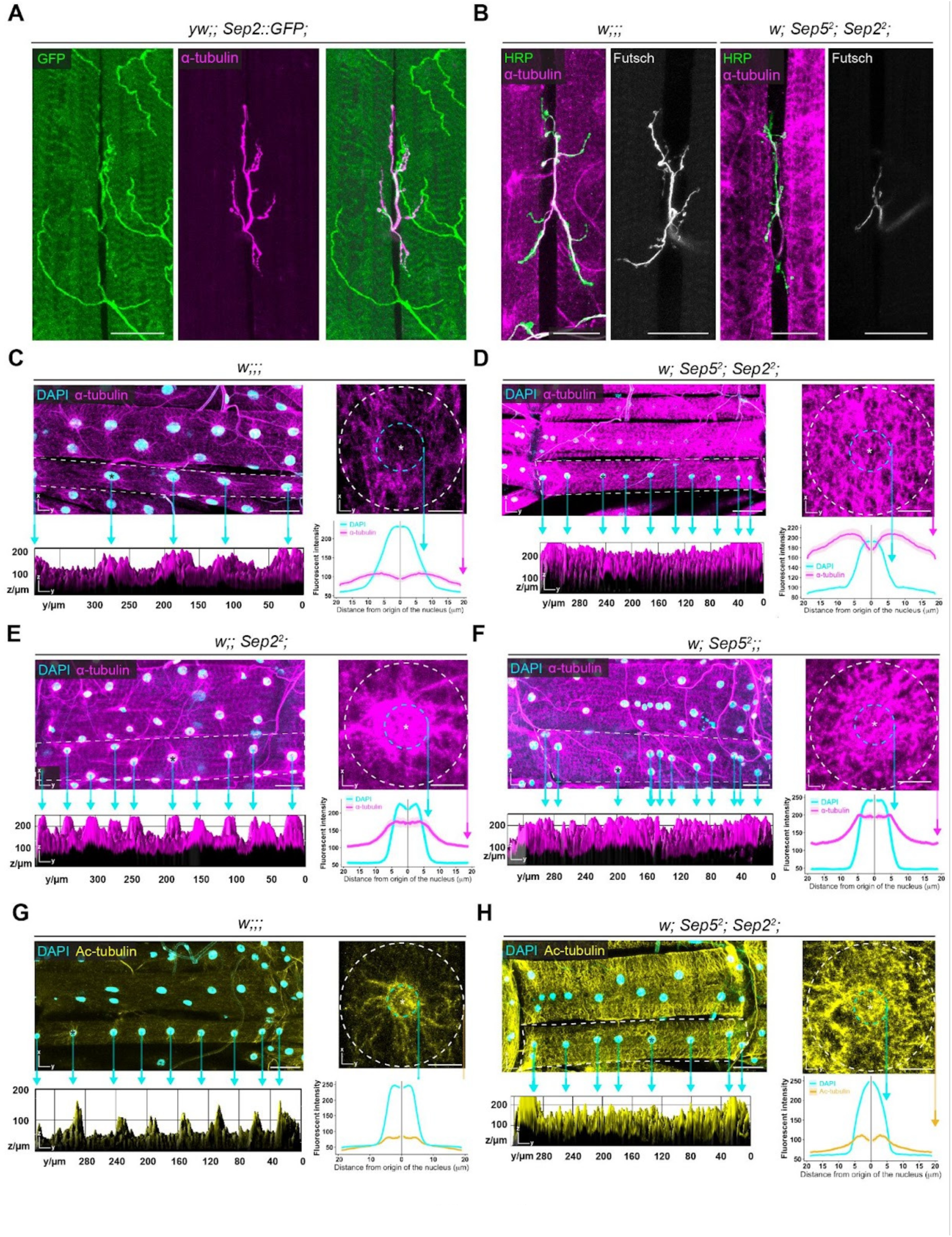
Septins spatially constrain microtubules along axons and around myonuclei and limit the accumulation of acetylated tubulin. (A) Confocal images of third-instar larval motor neurons expressing endogenously tagged Sep2::GFP (green) together with α-tubulin (magenta). Sep2::GFP localizes along axonal shafts and terminal branches and overlaps with α-tubulin, indicating that Sep2 associates with the microtubule cytoskeleton. (B) Representative images of motor axons at the larval neuromuscular junction labeled with anti-HRP (green), α-tubulin (magenta), and the microtubule-associated protein Futsch (white) in control (*w;;;*) and double mutant (*w; Sep5²; Sep2²*) larvae. In controls, Futsch-positive NMJ terminals form microtubule loops, whereas in double mutants, microtubule organization in NMJ terminals appears altered and has not formed well-structured loops. (C–H) Organization of microtubules in larval body wall muscle 6 (abdominal segment A4) stained for α-tubulin (magenta) or acetylated tubulin (Ac-tubulin, yellow) together with nuclei labeled by DAPI (cyan). In control muscles (C), microtubules are arranged in radial arrays with pronounced perinuclear enrichment. The white-dashed rectangle (C–H) indicates the regions used for 3D surface-plot analysis, with corresponding intensity profiles shown below. Radial intensity profiles of individual nuclei are shown on the right; cyan dashed circles mark the DAPI-defined nuclear boundaries, and white dashed circles indicate the region used for radial analysis (radius: 100 px). In double mutants (D), the α-tubulin signal is increased and redistributed, with loss of tight perinuclear confinement and elevated microtubule signal across the muscle fiber. (E–F) Microtubule organization in single mutant muscles from *Sep2²* (C) and *Sep5²* (D) larvae shows expanded perinuclear microtubule accumulation compared to controls, while largely preserving overall radial organization. (G–H) Distribution of stabilized microtubules visualized by acetylated tubulin staining in control (G) and double mutant (H) muscles. In controls, acetylated microtubules are enriched around myonuclei, whereas in double mutants, acetylated tubulin levels are increased and broadly distributed throughout the muscle. 3D surface-plot and radial intensity analyses reveal reduced spatial restriction of acetylated microtubules relative to nuclear boundaries. Data are presented as mean ± SEM. Scale bars: 40 µm (A, B, and whole-muscle views in C–H), 10 µm (single-nucleus views in C–H). Images were acquired by confocal microscopy using a 40X objective.

### The loss of Sep2 and Sep5 disrupts the formation of microtubule terminal loops at neuromuscular junctions (NMJs) and affects the organization of microtubules in larval muscle tissue

Furthermore, with regard to the expression and colocalization of septins with microtubules in motor neurons, attention was given to their presence on axonal branches within muscles and at NMJs.

To determine whether Sep2 and/or Sep5, along with their interaction with microtubules, are essential for proper microtubule organization and stabilization at NMJs, the expression pattern of Sep2::GFP was initially examined. It was observed that Sep2::GFP colocalized with microtubule loops at NMJ terminals and showed extensive colocalization in distal axonal branches (Fig. 1A).

To investigate the role of septins in microtubule organization, muscle 6/7 was stained using HRP and α-tubulin. Controls (*w;;;*), displayed clear labeling of motor axons and fine terminal branches by α-tubulin, with normal bouton-associated extensions (Fig. 1A). In contrast, analysis of *Sep2²*, *Sep5²* double mutants revealed minimal α-tubulin labeling in axonal terminal branches, indicating defective microtubule organization within the NMJ. Further assessment focused on Futsch, a microtubule-associated protein (MAP) that stabilizes microtubules and supports synaptic growth, acting as a linker between presynaptic microtubules and active zone components^38^. In control larvae, Futsch was well-organized along axonal projections, branches, and synaptic boutons, forming characteristic microtubule loops. Conversely, in double mutants, Futsch remained predominantly within the main axonal projection, with reduced branching and a loss of bouton-associated microtubule loops (Fig. 1B).

Additionally, we observed increased α-tubulin labeling in muscle tissue (Fig. 1B), suggesting dysregulation of tubulin levels or abnormalities in polymerization and localization. To investigate further, the organization of microtubules in larval body wall muscle 6 was analyzed through staining for α-tubulin and acetylated tubulin. In control muscles (Fig. 1C), microtubules displayed a radial arrangement with perinuclear enrichment, as demonstrated by 3D surface-plot and intensity profile analyses. In double mutants (Fig. 1D), α-tubulin signals were redistributed in the muscle area, while exhibiting a loss of clear perinuclear enrichment and elevated microtubule levels across the muscle fiber. Mutant muscles from *Sep2²* (Fig. 1E) and *Sep5²* (Fig. 1F) larvae showed increased microtubule intensity in the perinuclear region and across the muscle fibers while maintaining partial radial organization. Distribution analysis of acetylated tubulin, a marker for stabilized microtubules, revealed enrichment around myonuclei in control muscles (Fig. 1G). In contrast, double mutants exhibited increased levels of acetylated microtubules that were broadly dispersed throughout the muscle tissue. 3D surface plots and radial intensity analyses also demonstrated reduced spatial restriction of acetylated microtubules relative to nuclear boundaries (Fig. 1H).

Overall, these findings indicate that Sep2 and Sep5 are necessary for preserving presynaptic microtubule organization, as evidenced by α-tubulin and Futsch staining, and for maintaining microtubule organization in muscles, particularly in the perinuclear region. Our data further suggest that these septins play a role in microtubule polymerization and depolymerization. Together, these results underscore a broader role for septins in coordinating cytoskeletal architecture at NMJs and in muscle tissue.

### Loss of Sep2 and Sep5 disrupts myonuclear positioning

Considering the established association between microtubule architecture and muscle nuclei and with regard to observed nuclear positioning in double and single mutants, we analyzed the spatial distribution of myonuclei in whole larval muscles utilizing HRP, actin, and DAPI staining in larval muscle 6 (Supp Fig. 2). In controls, myonuclei were uniformly distributed along the muscle fibers and maintained a well-defined, rounded morphology. In contrast, muscles deficient in Sep2 and Sep5 displayed clustered nuclei with altered nuclear morphology, including changes in shape and size, along with irregular inter-nuclear spacing (Supp Fig. 2A). This phenotype is schematically summarized in Supp Fig. 2B. Quantitative analysis revealed a significant reduction in the minimum nearest-neighbor distance between nuclei (p < 0.001***, Supp Fig. 2C), and the overall distribution of inter-nuclear distances shifted toward lower values (p < 0.0001****, Supp Fig. 2D). While control muscles exhibit evenly spaced myonuclei along the muscle fiber, double mutants show irregular nuclear positioning and clustering.

Collectively, these findings demonstrate that Sep2 and Sep5 are essential for preserving the organization, placement, and morphology of myonuclei in *Drosophila* larval muscles. Furthermore, these results suggest a broader role for septins in coordinating cytoskeletal organization within muscle tissue.

### Loss of Sep2 and Sep5 disrupts NMJ morphology and postsynaptic organization

To examine whether Sep2 and/or Sep5, and their interaction with microtubules, are necessary for proper NMJ morphology, we analyzed neuromuscular junctions (NMJs) in muscles 6 and 7 within abdominal segment A4 of third-instar larvae using the null alleles *Sep2²* and *Sep5²*. Presynaptic membranes were labeled with HRP^39^, and postsynaptic scaffolds were visualized with DLG1, the *Drosophila* homolog of mammalian PSD-95, essential for synapse formation at embryonic NMJs^40^. Control NMJs displayed the typical bead-like bouton organization, with clear HRP-positive presynaptic outlines surrounded by DLG1-positive postsynaptic structures (Fig. 2A). Both *Sep2²* and *Sep5²* single mutants showed significant increases in branch number (p < 0.001 and p < 0.0001, respectively; Fig. 2C) and bouton number (p < 0.01 and p < 0.0001, respectively; Fig. 2D), indicating synaptic overgrowth while largely maintaining bouton identity. In contrast, *Sep2²*; *Sep5²* double mutants exhibited the most severe defects: HRP staining revealed extensive presynaptic membrane disorganization and loss of discrete bouton morphology, along with disrupted DLG1 distribution, indicating widespread structural disruption across the synapse (Fig. 2B). In these double mutants, bouton counts could not be reliably determined due to profound morphological disruption (Fig. 2E), and the number of branches was reduced compared to controls (Fig. 2F). To quantify changes in synaptic compartmentalization, we measured DLG1–HRP overlap as the fraction of HRP-positive presynaptic area that was also DLG1-positive. This analysis showed significant increases in overlap at Sep2², Sep5², and double-mutant NMJs (p < 0.01, p < 0.0001, and p < 0.0001, respectively; Fig. 2G), consistent with altered organization of pre- and postsynaptic domains. To further evaluate postsynaptic receptor patterning, we examined the distribution of the A-type glutamate receptor subunit GluRIIA. In controls, GluRIIA formed discrete, well-defined clusters at synaptic boutons (Fig. 2H). Conversely, double mutants showed increased receptor density, with GluRIIA clusters that were more dispersed and less confined to bouton regions (Fig. 2I). Collectively, these results demonstrate that Sep2 and Sep5 regulate NMJ growth, maintain bouton architecture, and ensure proper localization of postsynaptic DLG1 and GluRIIA-containing receptor fields. The combined loss of Sep2 and Sep5 causes severe synaptic disorganization and postsynaptic patterning defects.

**Figure 2.**
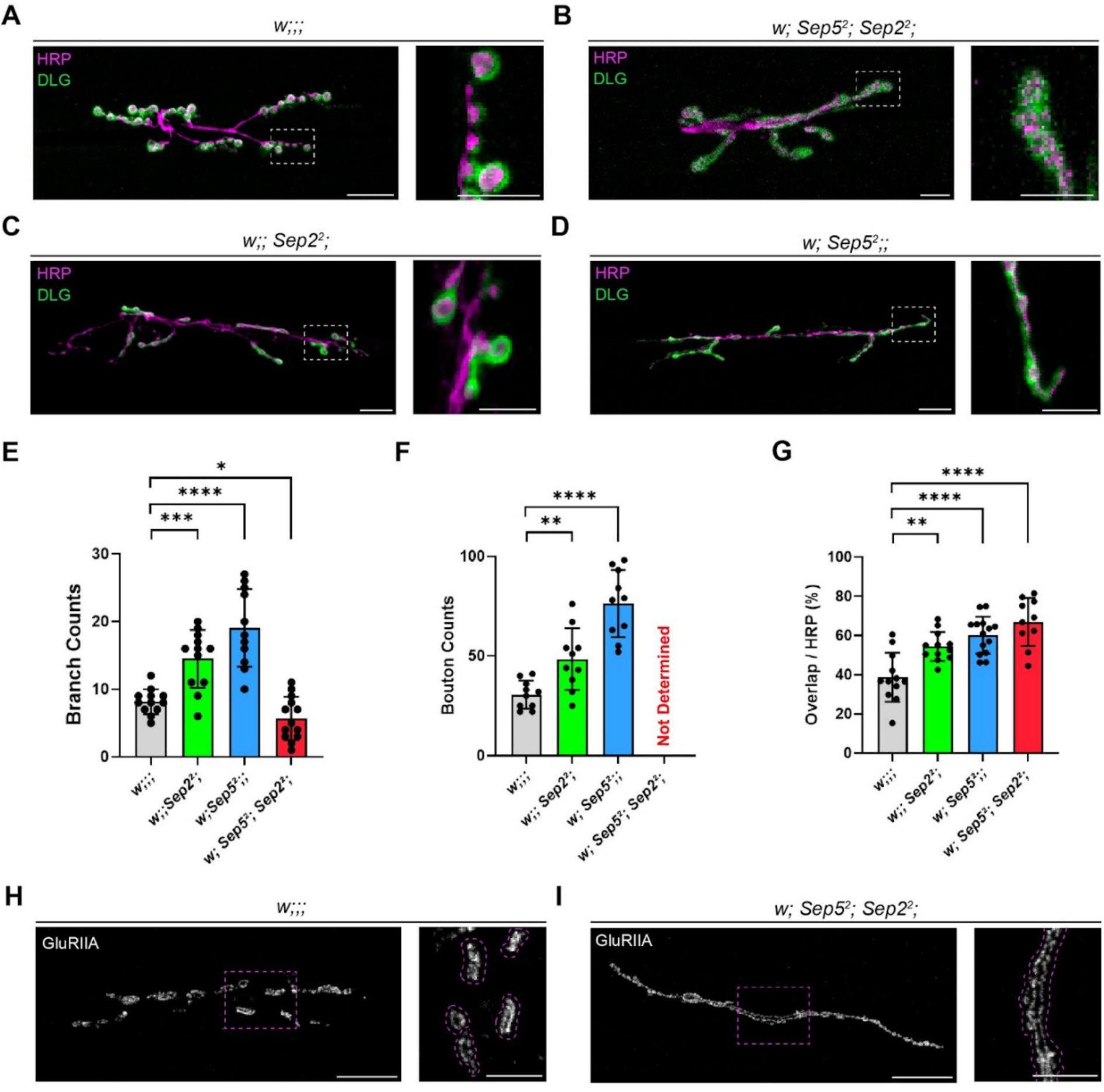
Sep2 and Sep5 are required for proper synaptic organization and postsynaptic receptor patterning at the larval neuromuscular junction. (A–D) Representative confocal images of neuromuscular junctions (NMJs) at muscle 6/7, abdominal segment A4, from third-instar larvae of the indicated genotypes. Presynaptic neuronal membranes are labeled with anti-HRP (green), and the postsynaptic density marker DLG is shown in magenta. Control NMJs (*w;;;*) display stereotypical, well-defined boutons with clear DLG rings surrounding HRP-positive presynaptic membranes (A). In double mutants (*w; Sep5²; Sep2²*), NMJs exhibit severe architectural disorganization, elongated structures, and loss of discrete bouton morphology (B). Sep2² (C) and Sep5² (D) single mutants show increased branching and bouton numbers compared to controls, while largely preserving overall bouton identity. Right panels show higher-magnification views of the dashed boxed regions. (E–F) Quantification of NMJ branch numbers (E) and bouton counts (F). Both Sep2² and Sep5² single mutants exhibit significantly increased branching and bouton numbers relative to controls. Bouton counts could not be determined for double mutants due to severe morphological disruption. (G) Quantification of DLG–HRP overlap, expressed as the percentage of HRP-positive areas overlapped by DLG. Double mutants show a significant increase in overlap compared to controls and single mutants, consistent with impaired distribution of pre- and postsynaptic compartments. (H–I) GluRIIA immunostaining of NMJs from control (H) and double mutant (I) larvae. Boxed regions are shown at higher magnification; dashed outlines indicate individual glutamate receptor clusters. In controls, GluRIIA clusters are compact and restricted, whereas in double mutants, they appear irregular and dispersed along the synaptic arbor. All images are maximum-intensity projections. Quantifications were performed in Fiji; n > 10 NMJs per genotype. Data are shown as mean ± SEM. Statistical analyses were performed using parametric tests after normality assessment (Shapiro–Wilk). Significance levels: ****p < 0.0001, ***p < 0.001, **p < 0.01, *p < 0.05. Scale bars: 20 μm (main panels), 10 μm (zoomed panels).

### Loss of septins disrupts presynaptic compartmentalization and active zone organization at the NMJ

To explore how septin depletion impacts presynaptic structure, we analyzed Synapsin (Syn) and Bruchpilot (BRP), which are crucial for synaptic vesicle clustering and active zone (AZ) structure^41^. In control larvae, Syn was localized within distinct puncta in boutons, overlapping with HRP-positive presynaptic membranes (Fig. 3A). Single mutants showed significantly increased Syn/HRP ratios (*Sep2²*: ns, *Sep5²*: p < 0.05, Fig. 3B), suggesting an expanded distribution of Synapsin. In double mutants, Syn localization was drastically altered (p < 0.0001): it appeared diffuse, spread throughout the NMJ arbor, and lacked the discrete bouton confinement seen in controls (Fig. 3A, B). BRP staining in controls revealed the expected punctate active zone pattern. In double mutants, BRP was mislocalized, losing its specific AZ distribution across the presynaptic membrane (n=14–18 NMJs per genotype; Fig. 3C). These results demonstrate that septins are essential for maintaining presynaptic compartmentalization of Syn and BRP and for preserving active zone organization.

**Figure 3.**
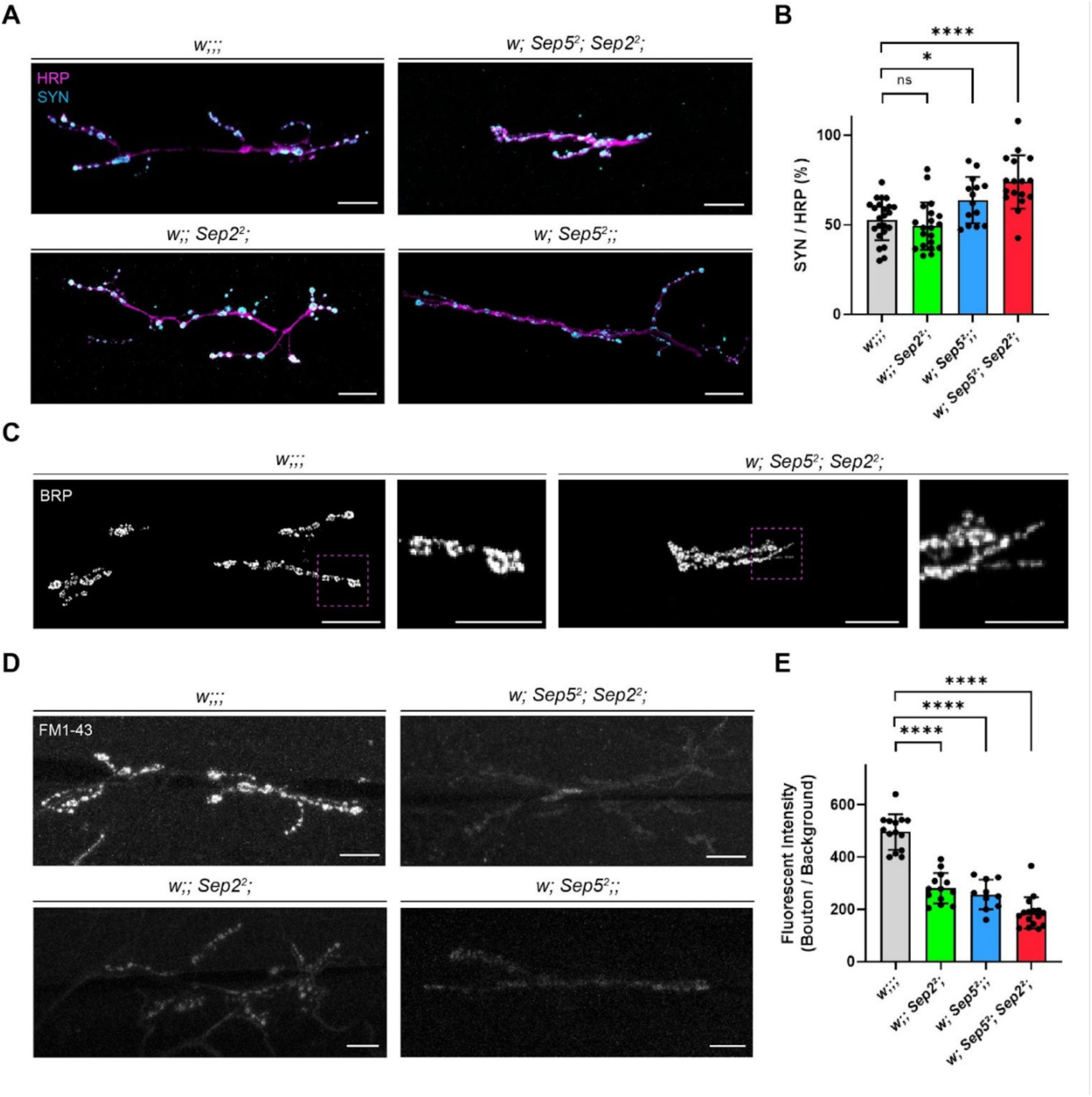
Septin loss alters presynaptic protein organization and impairs synaptic vesicle dye uptake at the *Drosophila* NMJ. (A) Representative confocal images of third-instar larval neuromuscular junctions (NMJs) at muscle 6/7, abdominal segment A4. Presynaptic neuronal membranes are labeled with anti-HRP (magenta), and synapsin (SYN; cyan) marks synaptic vesicle-enriched regions. Compared to controls (*w;;;*), *Sep2²* and *Sep5²* single mutants show a mildly increased SYN signal overlapping HRP-positive presynaptic membranes, whereas double mutants (*w; Sep5²; Sep2²;*) display visibly increased SYN-HRP colocalization, along with disrupted bouton morphology. (B) Quantification of SYN/HRP overlap, expressed as the percentage of HRP-positive areas overlapped by SYN. Double mutants show a significant increase compared to controls and single mutants (*Sep2²* vs control is not significant). (C) Bruchpilot (BRP) immunostaining (active zone marker) in control and double mutant NMJs. Boxed regions are shown at higher magnification. In controls, BRP puncta form discrete bouton-associated clusters, whereas in double mutants, BRP organization appears broadened and less sharply clustered into boutons. (D) FM1-43 dye uptake at NMJs following stimulation. In controls, FM1-43 labeling appears as a punctate bouton-associated signal, while all septin mutant genotypes exhibit reduced bouton FM1-43 intensity, with the lowest uptake in double mutants, where labeling is weak and diffuse. (E) Quantification of FM1-43 fluorescence intensity (bouton signal relative to background) reveals a significant reduction in septin mutants compared to controls. Quantifications were performed in Fiji; statistical analysis used Shapiro–Wilk normality testing followed by two-tailed t-tests for parametric datasets. Significance levels: ****p < 0.0001, *p < 0.05, ns: not significant. Scale bars: 20 μm (all panels).

### Loss of septins disrupts efficient synaptic vesicle recycling

Septin proteins have been linked to dynamin-dependent membrane processes, including endocytosis and vesicle transport^42^. Since dynamin (*Drosophila* shibire) is crucial for clathrin-mediated endocytosis at the NMJ^43^, we next tested synaptic vesicle recycling through FM1-43 dye uptake (Fig. 3D, E). After stimulation, control larvae showed strong FM1-43 uptake into presynaptic boutons, indicating effective vesicle recycling. In contrast, signal intensity was reduced in single mutants lacking Sep2 or Sep5 (Fig. 3D), implying partial impairment of vesicle endocytosis. Double mutants displayed markedly decreased FM1-43 labeling and nearly indiscernible bouton structures, revealing severe defects in synaptic organization as well as vesicle uptake and retention (Fig. 3D). Quantification of bouton/background fluorescence confirmed significant reductions in mutants, with the most severe defect observed in the double mutant (p < 0.0001, two-tailed t-tests) (Fig. 3E). These findings demonstrate that Sep2 and Sep5 are necessary not only for maintaining synaptic structure but also for efficient clathrin-dependent endocytosis and vesicle recycling. Alongside the Syn and BRP phenotypes, these results highlight an essential role for septins in organizing presynaptic architecture and function.

### Loss of Sep2 and Sep5 disrupts membrane organization and endocytosis beyond synapses

To better understand the role of septins in membrane dynamics and to assess whether septins are essential for membrane organization in highly secretory tissues, we examined third-instar larval salivary glands, which exhibit high levels of secretion, endocytosis, and vesicle trafficking. During larval handling and dissection, double mutants often showed melanization of the salivary glands, a phenotype not observed in controls or single mutants, prompting further investigation. Bright-field imaging revealed misshapen salivary glands with areas of tissue darkening in double mutants (Supp Fig. 3A). Immunostaining and confocal analysis demonstrated that, in controls, E-cadherin (E-cad) and DLG1 localized to apical/luminal junctions, outlining a distinct epithelial tube made up of properly shaped cells with evenly spaced, rounded nuclei (Supp Fig. 3B, C). In double mutants, the epithelium was severely disorganized. Cell borders were irregular, the lumen was dilated or misshapen, and both E-cad and DLG1 showed broadened and mislocalized junctional signals with increased intensity. Epithelial cells and their nuclei were enlarged and irregularly shaped. These results suggest that Sep2 and Sep5 are necessary to maintain junctional protein localization, epithelial structure, and lumen organization in the larval salivary gland. The observed phenotypes provide independent evidence of impaired membrane organization *in vivo* and support a broader role for septins in epithelial membrane dynamics and endocytosis beyond the nervous system.

### Loss of Sep2 and Sep5 causes locomotor defects in *Drosophila* larvae

Given our findings regarding defective synaptic structure, microtubule organization, and vesicle endocytosis, we asked whether neurotransmission deficits would translate into altered larval behavior. We recorded wandering third-instar larvae for two minutes under controlled environmental conditions and quantified their trajectories using FIMTrack software^44^ (n = 15–17 per genotype). See the methods and supplementary files for details on the experimental setup, behavioral data reproducibility, and data analysis. Control larvae displayed exploratory behavior with scattered, non-repetitive paths, whereas single mutants lacking Sep2 or Sep5 showed increased exploration. In contrast, double mutants moved in highly confined, often circular paths/trajectories (Fig. 4A). Quantitative analysis revealed that accumulated distance in double mutants was significantly reduced, and increased in single mutants as compared to controls (****p < 0.0001, Fig. 4C). Similarly, mean velocities (mm/s) were reduced in double mutants (****p < 0.0001, Fig. 4B). We next assessed movement persistence and posture (Fig. 4D-G). Double mutants spent less time in the Go-phase (percentage of frames moving), whereas single mutants displayed continuous crawling (Fig. 4D). Analyzed double mutant larvae also showed reduced time in a well-oriented, straight body configuration (head-tail axis: 180°; ****p < 0.0001; Fig. 4E). Consistent with this observation, double mutants frequently adopted a coiled posture (****p < 0.0001; Fig. 4H, I) and spent more time in both right- and left-bent states (****p < 0.0001; Fig. 4F, G). Together, these results demonstrate that the combined loss of Sep2 and Sep5 produces a robust locomotor phenotype, characterized by reduced distance and speed, shortened go-phase, defective body orientation with increased bending, and more frequent coiling. These behavioral impairments align with septin-dependent synaptic and cytoskeletal defects.

**Figure 4.**
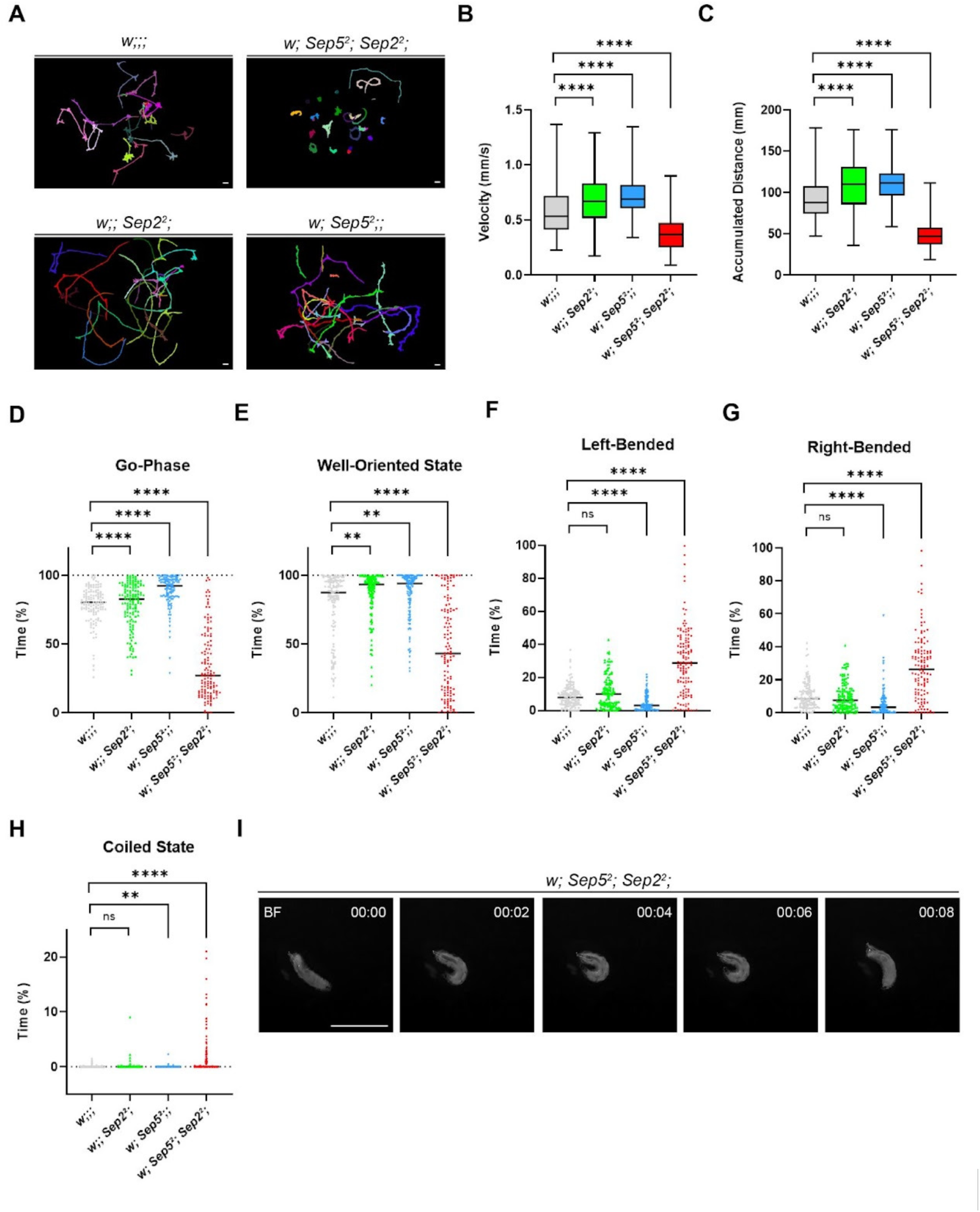
Septins are required to sustain coordinated locomotor state transitions during larval crawling. (A) Representative locomotion trajectories of individual third-instar larvae recorded over 120 s. Each colored trace represents a single larva. Control larvae (*w;;;*) display exploratory crawling with dispersed trajectories, whereas *Sep2²* and *Sep5²* single mutants exhibit continuous but less exploratory movement patterns. In contrast, double mutants (*w; Sep5²; Sep2²*) initiate movement but fail to sustain directional crawling, resulting in short, circular, or highly constrained trajectories. (B–C) Quantification of average crawling velocity (mm/s) (B) and accumulated distance traveled (mm) (C) across the full recording period. *Sep2²* and *Sep5²* single mutants show increased velocity and accumulated distance relative to controls, whereas double mutants exhibit a marked reduction in both parameters. (D–H) Behavioral state analysis based on frame-by-frame classification of larval posture and movement. The percentage of frames in which larvae occupied each behavioral state during the 120 s recording is shown; each dot represents an individual larva, and horizontal lines indicate the mean. (D) Double mutants spend significantly less time in the go-phase, defined as frames in which larvae actively propel themselves forward. (E) The well-oriented state, defined by proper alignment of anterior, middle, and posterior body segments, is reduced in double mutants but increased in single mutants. (F–G) Double mutants show a significant increase in both left- and right-bended states, indicating impaired motor coordination, without directional bias. (H) Time spent in the coiled state is significantly elevated in double mutants, reflecting failure to maintain coordinated body posture. (I) Representative bright-field (BF) time series showing a double mutant larva transitioning into a coiled, stuck motor state, characterized by loss of posterior propagation and body alignment. Statistical analyses were performed following Shapiro–Wilk normality testing, using two-tailed t-tests for parametric and Mann–Whitney tests for nonparametric datasets. Asterisks denote statistical significance (****p < 0.0001, ***p < 0.001, **p < 0.01, *p < 0.05; ns, not significant). Sample sizes: *w;;;* (n = 129), *w;; Sep2²;* (n = 133), *w; Sep5²;;* (n = 118), *w; Sep5²; Sep2²;* (n = 126). Scale bars: 4 mm.

### Septin Reconstitution Partially Restores Cytoskeletal Organization and Synaptic Function in Double Mutants

To determine whether the microtubule disorganization, endocytic defects, and behavioral abnormalities observed in *Sep2²* and Sep5² double mutants result directly from septin loss, we generated a genetic rescue in which Sep2::GFP is expressed under its endogenous promoter in the double-mutant background (*w; Sep5²; Sep2², Sep2::GFP*). Biallelic endogenous expression restored viability: whereas *Sep2²; Sep5²* larvae arrested at the early pupal stage, rescue animals developed into viable adults, demonstrating functional complementation by Sep2::GFP.

At larval neuromuscular junctions (NMJs), Sep2::GFP colocalized with HRP-positive presynaptic membranes (Fig. 5A), confirming correct axonal targeting in the double-mutant background and consistent localization with that observed in non-mutant animals (Fig. 1A, Supp Fig. 1).

**Figure 5.**
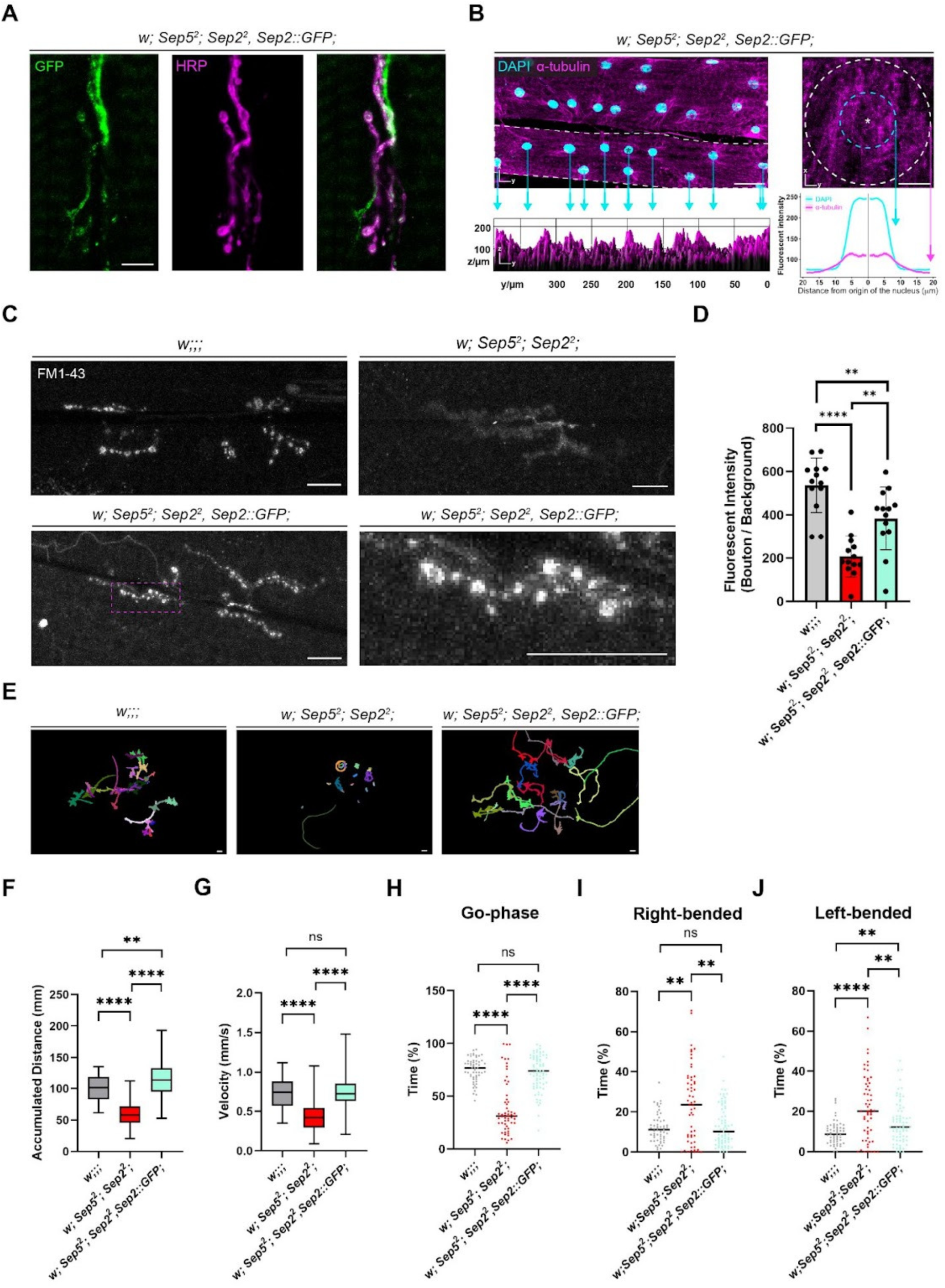
Endogenously expressed Sep2::GFP partially rescues synaptic vesicle uptake and locomotor coordination defects in Sep2 and Sep5 double mutants. (A) Representative confocal images of larval NMJs showing Sep2::GFP signal (green) in the *w; Sep5²; Sep2², Sep2::GFP* background (*Sep2::GFP* cassette driven by the endogenous promoter). Presynaptic membranes are labeled with anti-HRP (magenta). Note the region where Sep2::GFP exhibits a bouton-associated pattern aligned with HRP-positive presynaptic structures (B). Larval body wall muscle 6 (abdominal segment A4) stained for α-tubulin (magenta) and nuclei (DAPI, cyan) in the double mutant rescue (*w; Sep5²; Sep2², Sep2::GFP)* background. Radial intensity analysis centered on an individual nucleus is shown to visualize perinuclear microtubule distribution. Sep2::GFP provides partial rescue of microtubule organization, while nuclear localization remains incompletely restored. (C) FM1-43 uptake at NMJs following stimulation. Control (*w;;;*) boutons show punctate FM1-43-labeled vesicle clusters, whereas double mutants (*w; Sep5²; Sep2²*) show reduced and diffuse labeling. In double mutant rescue (*w; Sep5²; Sep2², Sep2::GFP)*, punctate bouton-associated FM1-43 signal and overall intensities are restored toward control levels. The dashed magenta box indicates the region shown at higher magnification. (D) Quantification of FM1-43 fluorescence intensity (bouton signal relative to background) shows reduced uptake in double mutants and significant improvement in double mutant rescue animals. (E) Representative locomotion trajectories of individual third-instar larvae recorded over 120 s; each colored trace represents a single larva. Double mutants exhibit highly constrained movement, whereas double mutant rescue *(w; Sep5²; Sep2², Sep2::GFP)* larvae display increased exploratory crawling compared to double mutants, approaching control behavior. (F–G) Quantification of accumulated distance traveled (F) and average velocity (G) across the recording period. double mutant rescue animals (*w; Sep5²; Sep2², Sep2::GFP)* move significantly more than double mutants and trend toward control values. (H–J) Behavioral state analysis showing the percentage of frames spent in the go-phase (H), right-bended (I), and left-bended (J) states during the recording. Double mutants spend less time in the go-phase and more time in bending states, whereas double mutant rescue larvae (*w; Sep5²; Sep2², Sep2::GFP)* shift these measures toward control levels, indicating improved motor coordination. Statistical analyses were performed following Shapiro–Wilk normality testing, using two-tailed t-tests for parametric and Mann–Whitney tests for nonparametric datasets. Asterisks denote statistical significance (****p < 0.0001, **p < 0.01; ns, not significant). Scale bars: 10 μm (A and single-nucleus views in B), 40 µm (B whole-muscle views), 4mm (E).

In larval body wall muscle 6 (A4), α-tubulin staining combined with radial intensity profiling and 3D surface analysis revealed partial restoration of perinuclear microtubule organization. Microtubule distribution shifted toward a radial, control-like arrangement with partial re-establishment of perinuclear organization (Fig. 5B); however, nuclear positioning defects were not fully corrected, indicating incomplete rescue of the muscle phenotype.

To assess synaptic vesicle recycling, we performed FM1-43 uptake assays following stimulation. Double mutants exhibited diffuse and markedly reduced bouton-associated fluorescence, consistent with impaired endocytosis. In contrast, rescue animals exhibited restored punctate FM1-43 labeling (Fig. 5C) and significantly higher fluorescence intensity than double mutants (Fig. 5D), approaching control levels, indicating partial recovery of synaptic vesicle endocytosis.

Locomotor tracking of third-instar larvae over 120 s further demonstrated that rescue animals exhibited increased exploratory crawling compared to double mutants (Fig. 5E). Accumulated distance traveled (Fig. 5F) and average velocity (Fig. 5G) were significantly elevated and trended toward control values. Behavioral state analysis revealed increased go-phase occupancy (Fig. 5H) and reduced bending states (Fig. 5I, 5J), consistent with improved motor coordination.

Collectively, these findings demonstrate that endogenous Sep2 expression partially restores viability, cytoskeletal organization, synaptic vesicle recycling, and coordinated locomotion in double mutants, supporting a model in which septins function as critical organizers that link cytoskeletal and microtubular architecture to presynaptic membrane dynamics and motor output.

### Allele-Specific Effects of Sep2 Reveal Cytoskeletal and Functional Consequences at the NMJ

To test whether septin phenotypes depend on allele class, we analyzed a second Sep2 mutant^3^, Sep2¹ (Δ∼1.1 kb). We predict that Sep2¹ retains an intact GTP-binding G-domain, allowing its incorporation into septin filaments. Sep5 compensation is unlikely to be required in the Sep2¹ background.

In *Drosophila melanogaster*, septin protofilaments assemble as hexameric complexes. Within this structure, the G-interface mediates intermolecular interactions between septin subunits, particularly within the Sep1–Sep2 heterodimer and Pnut–Pnut homodimers. Proper G-interface interactions are essential for stable heterodimer formation and higher-order filament assembly^2,45^.

AF2-predicted structures indicate that Sep2¹ retains an intact portion of the G-domain, suggesting the potential to remain structurally compatible with incorporation into septin protofilaments, in contrast to Sep2², which appears to disrupt septin architecture (Fig. 6A). AF2 model confidence and agreement with the Sep1–Sep2 crystal structure (PDB: 8DKT) were assessed by pLDDT and PAE across wild-type Sep2 models (Sep2RA, Sep2RB) and mutant variants (Sep2¹, Sep2²). Overall, the models showed high local confidence (average pLDDT > 80), with 42.1% of residues in the very high-confidence range (predominantly within the G-domain) and 39.1% in the high-confidence range (encompassing switch regions, trans-loops, and the C-terminal segment). While the G-domain appears compact and well defined (PAE < 5 Å), the relative placement of the G-domain with respect to the C-terminal segment is uncertain, with predicted errors reaching ∼30 Å. Consistent with these predictions, an over 600 ns MD simulation of the Sep1–Sep2¹ complex supported the potential for Sep2¹ incorporation via a distorted canonical interface motif. Although level III interactions mediated by switch regions and trans-loops were missing, Sep2¹ remained bound to Sep1 through reorganized level I and level II contacts (Supp. Fig. 4). Details of the MD simulation and trajectory analysis are provided in the Methods and Supplementary Files. Collectively, these analyses suggest that Sep2¹ is structurally competent yet functionally altered as a septin subunit.

**Figure 6.**
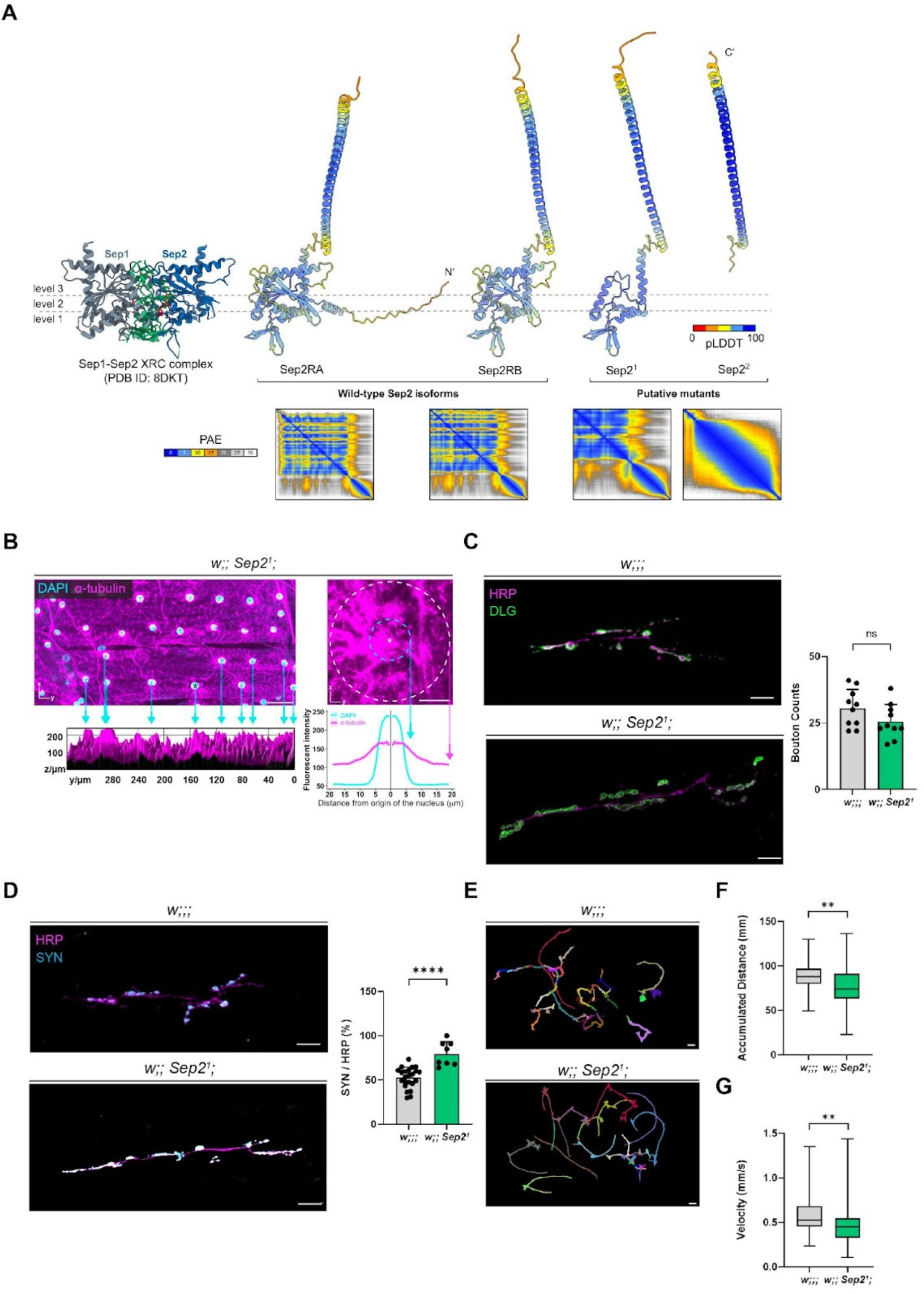
The Sep2¹ allele perturbs microtubule organization, alters presynaptic protein distribution, and impairs larval locomotion. (A) The X-ray crystal structure (XRC) of the Sep1–Sep2 complex (PDB ID: 8DKT) is shown alongside AF2-predicted models of wild-type Sep2 (Sep2RA, Sep2RB) and putative mutants Sep2 (Sep2^1^, Sep2^2^). AF2 structures are colored by per-residue pLDDT confidence (0–100), and the corresponding PAE matrices (0–30 Å) are shown for each structure. Sep1–Sep2 interface contacts are evaluated at three levels (I–III), with interface residues highlighted in green and GDP/GTP in orange. (B) Organization of microtubules in larval body wall muscle 6 (abdominal segment A4) stained for α-tubulin (magenta) and nuclei (DAPI, cyan) in control (*w;;;*) and *Sep2¹* mutant (*w;; Sep2¹;*) larvae. In *Sep2¹* mutants, α-tubulin signal shows broadened perinuclear enrichment and altered spatial distribution compared to controls. Cyan arrows indicate positions used for 3D surface analyses, with corresponding intensity profiles shown below. Radial intensity profiles centered on individual nuclei are shown on the right; cyan dashed circles mark the DAPI-defined nuclear boundaries. (C) Representative confocal images of neuromuscular junctions (NMJs) labeled with anti-HRP (magenta; presynaptic membrane) and DLG (green; postsynaptic density marker) in control and *Sep2¹* mutant larvae. Quantification of bouton number shows no significant difference between genotypes (right panel), indicating preserved gross NMJ morphology. (D) Representative images of NMJs stained for HRP (magenta) and synapsin (SYN; cyan). *Sep2¹* mutants display increased SYN localization relative to HRP compared to controls. Quantification of SYN/HRP overlap shows a significant increase in *Sep2¹* mutants, indicating altered presynaptic vesicle organization. (E) Representative locomotion trajectories of individual third-instar larvae recorded over 120 s. Control larvae exhibit exploratory crawling with dispersed trajectories, whereas *Sep2¹* mutants display shorter and more constrained movement paths. (F-G) Quantification of accumulated distance traveled (F) and average crawling velocity (G) during the recording period. *Sep2¹* mutants travel significantly shorter distances and move more slowly than controls. Box plots show the median (horizontal line) and distribution of values. Statistical comparisons were performed using Shapiro–Wilk normality testing followed by two-tailed t-tests for parametric data and Mann–Whitney U tests for nonparametric distributions. p < 0.01; ns, not significant. Scale bars: 10 μm (single-nucleus views in B), 20 μm (C-D), 40 µm (B whole-muscle views), 4mm (E).

At the tissue level, analysis of larval muscle 6 stained for α-tubulin and DAPI revealed enhanced perinuclear α-tubulin enrichment and altered radial microtubule distribution in Sep2¹ mutants, supported by radial intensity profiling and 3D surface analyses (Fig. 6B). Overall α-tubulin intensity was increased compared to controls, a phenotype consistent with other septin mutants. Thus, incorporation of a structurally compatible but mutant Sep2 subunit is sufficient to perturb cytoskeletal organization.

At the neuromuscular junction (NMJ), gross synaptic architecture was preserved in Sep2¹ mutants. Quantification of bouton number revealed no significant difference between genotypes (Fig. 6C). Representative NMJ images stained for HRP and Synapsin (SYN) showed increased SYN localization relative to HRP in Sep2¹ mutants compared to controls (Fig. 6D). Quantification of SYN/HRP overlap demonstrated a significant increase in Sep2¹ mutants, consistent with altered presynaptic vesicle organization despite preserved bouton number.

Functionally, Sep2¹ larvae exhibited shorter and severely restricted locomotor trajectories than controls (Fig. 6E). Accumulated distance traveled (Fig. 6F) and average velocity (Fig. 6G) were significantly reduced (p < 0.01), and Go-phase occupancy was decreased (Supp. Fig. 5A), whereas orientation (Supp. Fig. 5B), bending (Supp. Fig. 5D, 5E), and coiling (Supp. Fig. 5C) parameters were unchanged.

Collectively, these findings demonstrate that incorporation of a mutant but assembly-competent Sep2 subunit disrupts microtubule organization and presynaptic vesicle distribution without altering bouton number. In contrast to Sep2², which compromises septin architecture and synaptic structure, Sep2¹ selectively perturbs filament function. Thus, septin loss and septin misincorporation impair NMJ function through distinct but convergent cellular mechanisms.

### Combined Loss of Sep2 and Sep5 Drives Stabilization-Dominant Cytoskeletal Transcriptional Reprogramming

Septins are GTP-binding scaffolds that regulate kinesin- and dynein-mediated transport by controlling motor–cargo interactions on microtubules. Through specific paralog and isoform combinations, they coordinate vesicular trafficking and membrane dynamics^14^ .Alterations in their higher-order architecture can reorganize the cytoskeleton and modify trafficking-dependent signaling, thereby indirectly shaping transcriptional programs and gene expression.

To obtain an unbiased organism-wide view of the pathways perturbed by combined Sep2/Sep5 loss and to distinguish double-mutant effects from potential compensation in single mutants, we performed RNA-seq on wandering L3 male larvae, to directly align with the developmental stage used for neuromuscular junction (NMJ) and behavioral assays. Whole-larva profiling captured both neuronal and non-neuronal tissues and generated a comprehensive atlas of transcriptional changes and candidate genes for targeted validation and genetic testing.

Principal component analysis (PCA) of TMM-normalized log-CPM values across biological replicates (Fig. 7A) revealed strong genotype-dependent clustering with tight grouping of replicates, confirming reproducibility and minimal batch contribution. PC1 accounted for 44.1% of total variance and robustly separated *Sep2²*, *Sep5²* double mutants (SepDM) from wild-type and both single mutants, identifying combined septin loss as the dominant driver of global transcriptional variation. PC2 explained 11.5% of the variance and segregated Sep2 single mutants along an independent axis, indicating genotype-specific transcriptional effects. These data demonstrate that *Sep2²*, *Sep5²* double mutants occupy a distinct transcriptional space that cannot be explained by additive single-mutant effects.

**Figure 7.**
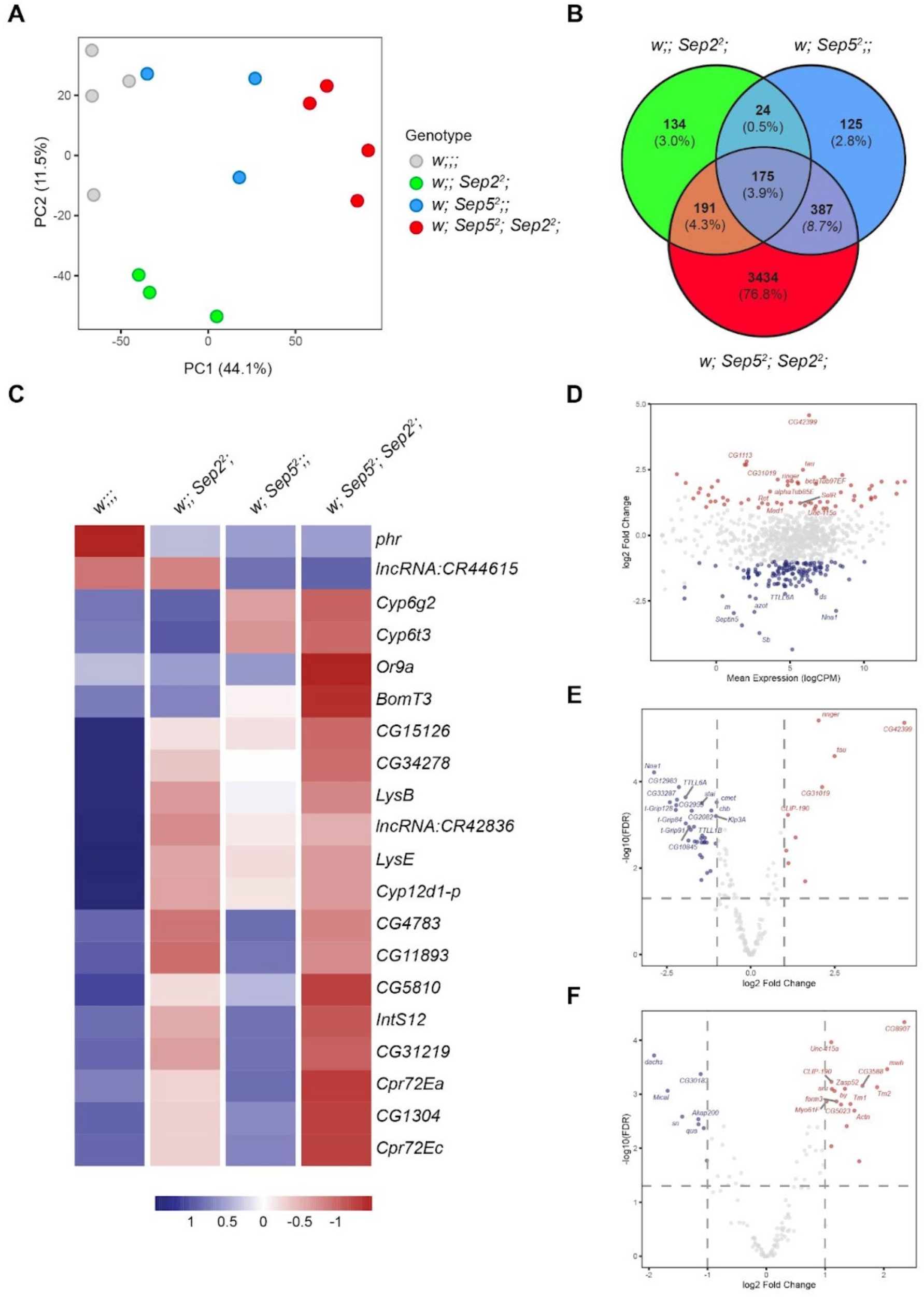
Transcriptomic disruption of the cytoskeletal network in *Sep2* and *Sep5* double mutants. (A) Global Principal Component Analysis (PCA): Multi-dimensional scaling of TMM-normalized log-CPM values across all biological replicates (n=3-4 per genotype). Samples are colored by genotype: *w;;;* (gray), *w;; Sep2*^2^; (green), *w; Sep5*^2^;; (blue), and the double mutant *w; Sep5²; Sep2²;,* (red). (B) Venn diagram illustrating the distribution of significant DEGs (FDR < 0.05, |log2 fold change| > 1) across the three mutant genotypes compared to the Wild-type (W) baseline. Circles are colored by genotype: *w; Sep5²; Sep2²;* (red), *w; Sep5*^2^;; (blue), and *w;; Sep2*^2^; (green). (C) Signature of Top 20 Differentially Expressed Genes (DEGs): Heatmap showing row-normalized Z-scores of the 20 most significant DEGs identified in the *w; Sep5²; Sep2²;* vs. *w;;;* comparison. Values represent the average expression across replicates for each genotype. The color scale ranges from blue (downregulated) to orange (upregulated), highlighting the unique and intensified expression signature of the double mutant. (D) Global Cytoskeletal Disruption (MA Plot): Scatter plot showing the relationship between mean expression (logCPM) and log2 fold-change (logFC) for genes associated with the cytoskeleton and cytoskeleton organization (GO:0005856, GO:0007010). Genes with significantly altered expression (FDR < 0.05) in *w; Sep5²; Sep2²;* vs. *w;;;* are highlighted in blue and orange. The top 20 most significant cytoskeletal regulators are labeled by symbol. E-F. Specific Vulnerability of Tubulin- and Actin-Binding Networks: Volcano plots illustrating the differential expression of genes involved in (E) Tubulin-binding (GO:0015631) and (F) Actin-binding (GO:0003779) in *w; Sep5²; Sep2²;* mutants. The y-axis represents statistical significance (-log10 FDR) and the x-axis represents the magnitude of change (log2 fold change). Dots are colored by regulation status: blue (downregulated, FDR < 0.05, logFC < -1), orange (upregulated, FDR < 0.05, logFC > 1), and gray (not significant). The top 20 most significant genes in each specific sub-category are labeled.

Differential expression analysis (FDR < 0.01; |log2FC| > 2) revealed a dramatic expansion of transcriptional perturbation uniquely in the double mutant (Fig. 7B). Sep2 and Sep5 single mutants exhibited 109 and 73 unique differentially expressed genes (DEGs), respectively, whereas *Sep2²*; *Sep5²* double mutants displayed 1,988 unique DEGs. Only 113 genes were shared across all genotypes. The magnitude and asymmetry of this expansion indicate a non-linear transcriptional response upon combined septin loss and support a redundancy/buffering model for Sep2 and Sep5 function.

Heatmap visualization of the top 20 most significant DEGs (Fig. 7C) in *Sep2²*; *Sep5²* double mutants versus wild-type revealed a coherent and intensified transcriptional signature specific to the double mutant. Several genes exhibited markedly elevated expression relative to both wild-type and single mutants, reinforcing that the double mutant state reflects synergistic transcriptional reprogramming rather than cumulative additive effects.

Gene Ontology enrichment analysis (Supp Fig. 6 A-F) further defined the directional nature of this remodeling. Up-regulated DEGs were enriched for peptidase/serine-hydrolase and oxidoreductase activities and for extracellular-matrix/external encapsulating structure terms. Down-regulated DEGs were enriched for phosphatase/phosphoric-ester–hydrolase activities and tubulin/microtubule binding, mapping to the microtubule cytoskeleton, cilia/axoneme, and the microtubule-organizing center. At the biological process level, metabolic and catabolic pathways were enriched among up-regulated genes, whereas cilium organization, microtubule-based movement, and plasma-membrane–bound cell-projection assembly were enriched among down-regulated genes. Together, Sep2/Sep5 loss suppresses microtubule-related modules and membrane–cytoskeletal dynamics while inducing metabolic-stress programs consistent with autophagy.

MA plot analysis (Fig. 7D) demonstrated widespread, yet structured, differential expression across a broad range of transcript abundance for cytoskeletal-related genes in Sep2²; Sep5² double mutants, excluding low-expression bias as the primary explanation. Refined volcano plot analysis identified significant dysregulation within tubulin-binding (Fig. 7E) and actin-binding (Fig. 7F) gene subsets, confirming that cytoskeletal networks are centrally and statistically robustly affected.

Targeted interrogation of cytoskeletal gene families (Supp data, Table 1) revealed a highly directional and asymmetric transcriptional remodeling in the Sep2²; Sep5² double mutant. Within microtubule-associated proteins (MAPs) and stabilizers, expression changes were selective rather than uniform. Among MAPs, tau (CG45110) exhibited the strongest induction (log2FC ≈ +2.33; ∼5-fold), representing the most prominent transcriptional change in this category. ringer (CG45057) was also robustly upregulated (log2FC ≈ +2.0), consistent with increased microtubule bundling capacity. In contrast, CLIP-190 (CG5020) showed more moderate induction (log2FC ≈ +1.01), while other microtubule-binding proteins displayed comparatively modest or heterogeneous shifts. This pattern indicates that septin loss preferentially enhances a subset of lattice- and bundling-associated stabilizers rather than broadly elevating all MAPs.

Importantly, the major regulators of tubulin acetylation, HDAC6 (CG6170) and Atat (αTAT) (CG3967), showed no significant transcriptional change, indicating that the previously observed increase in microtubule acetylation is unlikely to arise from altered expression of the core acetylation machinery. Instead, acetylation changes likely reflect secondary stabilization of long-lived microtubules.

In contrast to MAP induction, genes associated with microtubule nucleation and dynamic remodeling were predominantly downregulated. Multiple γ-tubulin ring complex components, including t-Grip84 (CG7716), t-Grip128 (CG32232), and t-Grip91 (CG18109), were significantly reduced (log2FC ≈ –1.9 to –2.2), arguing against increased nucleation capacity. Similarly, several kinesin family members, Klp59C, Klp59D, Klp3A, and Klp67A, were downregulated (log2FC ≈ –1.0 to –1.5), indicating reduced motor-driven transport potential rather than enhanced trafficking.

Consistent with reduced microtubule plasticity, enzymes that regulate post-translational modification of tubulin were also suppressed. Multiple TTLL family members (TTLL6A, TTLL1A, TTLL1B, TTLL4A, TTLL4B) exhibited significant downregulation, and the deglutamylase Nna1 was strongly reduced (log2FC ≈ –2.7). This coordinated decrease suggests diminished regulation of tubulin glutamylation, potentially affecting motor processivity and microtubule dynamics.

Beyond microtubules, actin-associated regulators showed bidirectional remodeling: mwh(CG43772) was significantly upregulated (log2FC ≈ +1.98), whereas dachs(CG42840) was markedly downregulated (log2FC ≈ –2.11). The extracellular remodeling enzyme Mmp2(CG1794) was also significantly reduced (log2FC ≈ –2.05), indicating coordinated alterations across cytoskeletal–membrane and extracellular interfaces.

Collectively, across stabilizers, nucleation factors, motors, and modification enzymes, the transcriptional landscape is neither globally suppressed nor uniformly activated. Instead, Sep2 and Sep5 loss produce a selective cytoskeletal reprogramming characterized by strong induction of lattice- and bundling-associated stabilizers (most prominently tau and ringer), coupled with suppression of nucleation machinery, motor systems, and post-translational modification enzymes. This pronounced asymmetry supports a stabilization-dominant compensatory state that favors the persistence of existing microtubule arrays over dynamic turnover, rather than generalized cytoskeletal collapse.

## Discussion

### Septins as Higher-Order Cytoskeletal Regulators

Septins are increasingly recognized as higher-order organizers of neuronal and synaptic architecture by coordinating microtubules, actin filaments, and membrane domains into integrated networks. Acting as membrane-associated scaffolds and diffusion barriers, septins have been implicated in regulating endocytosis, exocytosis, membrane remodeling, and actin dynamics^15,34,46–48^. In neurons, they associate with key proteins involved in vesicle trafficking and cytoskeletal rearrangement, influencing neurotransmitter release and vesicle positioning^22,49–51^. Through these scaffolding and compartmentalizing functions, septins exert spatial control over cytoskeletal dynamics, preserving neuronal integrity^25^. Dysregulation of septins has been associated with demyelinating, neurodevelopmental, neurodegenerative, and autoimmune disorders^25^.

### Septin Function in *Drosophila* NMJ

Using the genetically simplified septin system of *Drosophila*, we demonstrate that Sep2 and Sep5 act together to regulate microtubule stabilization and synaptic structural organization at the NMJ. Loss of these septins results in altered synaptic organization, impaired vesicle recycling, increased acetylated tubulin, and enhanced microtubule stabilization. By possibly constraining excessive stabilization, septins act as one of the key modulators of microtubule homeostasis. Septin loss is accompanied by a transcriptional shift consistent with enhanced microtubule stability. Thus, these findings position septins as regulators of microtubule state rather than simple structural stabilizers of synapses.

We show that both Sep2::GFP and Pnut display a highly organized and non-uniform distribution in Drosophila neurons, characterized by punctate assemblies within the soma and colocalization with α-tubulin along the axonal shaft. confirming that the tagged f Sep2::GFP is incorporated into septin filaments and directly associated with microtubules. Loss of both Sep2 and Sep5 disrupts bouton morphology, alters presynaptic compartmentalization of Synapsin and Bruchpilot, and perturbs postsynaptic regions (DLG1, GluRIIA). These structural alterations are accompanied by vesicle recycling defects, linking septin-dependent cytoskeletal architecture to presynaptic membrane dynamics.

These findings are consistent with previous mammalian studies showing presynaptic localization of Sept3 and Sept5, as well as NMJ enrichment of Sept9 and Sept5^19,51–53^, highlighting a conserved role for septins in presynaptic compartments. Sept5 has been shown to inhibit vesicle exocytosis by forming a structural barrier and modulating SNARE-mediated fusion, whereas Sept7 stabilizes dendritic spines and postsynaptic densities^54,55^.

At the *Drosophila* NMJ, postsynaptic PDZ-domain scaffolds such as DLG align glutamate receptors and cadherins with presynaptic specializations during synapse formation^56–58^. The balance between synapse stability and plasticity requires tight integration of signaling pathways with actin and microtubule networks^59^. Disruption of the presynaptic microtubule cytoskeleton is strongly associated with synapse destabilization and can promote synaptic disassembly, likely by weakening trans-synaptic protein interactions^60^.

Loss of Sep2 or Sep5 alters NMJ morphology, increasing bouton number and producing abnormal synaptic structures, while combined loss results in pronounced overgrowth with a disorganized bead-like bouton arrangement. Because Sep2 and Sep5 are required for assembly of the hexameric septin complex and incorporation of Pnut and Sep1/Sep4, their simultaneous absence likely abolishes compensatory complex formation, leading to severe cytoskeletal disorganization. Collectively, these results support a model in which septins maintain synaptic stability by coordinating actin and microtubule architecture, thereby preserving membrane integrity and proper bouton organization.

### Septin-Mediated Microtubule Organization

Septin-dependent microtubule regulation is not restricted to neurons. In larval muscles, Sep2, Sep5 double mutants exhibit broad microtubule disruptions: perinuclear enrichment of microtubules is lost, total α-tubulin signal is elevated around and between myonuclei, accompanied by a marked increase in α-tubulin acetylation. These observations align with mammalian Sept7 studies, where depletion leads to hyperacetylation, hyperstabilization, and impaired neuritogenesis due to reduced HDAC6-mediated deacetylation^12^.

Alongside the pronounced microtubule disorganization observed in muscle cells, we identified clustered myonuclei with irregular spacing and altered morphology in double mutants. These defects are consistent with established roles of the cytoskeleton in maintaining nuclear positioning and overall cytoskeletal architecture^61–63^.

Similarly, in highly secretory larval salivary glands, where sustained exocytosis must be counterbalanced by clathrin-mediated endocytosis to preserve apical membrane surface area^64^, Sep2 and Sep5 double mutants displayed disrupted, mislocalized junctional proteins, and enlarged and malformed lumen. Given that impaired endocytosis may lead to apical membrane overgrowth and lumen expansion^66^, these phenotypes suggest that septin complexes contribute to membrane turnover and apical surface restriction. Consistently, mutant glands display thickened, serrated apical membranes reminiscent of microvillar disorganization reported upon disruption of cytoskeletal-membrane coupling^66,68^. Together, these observations indicate that septins may spatially coordinate cytoskeletal architecture with membrane trafficking to maintain epithelial surface homeostasis.

The consistency of structural defects across neuronal and non-neuronal tissues argues for a conserved role of septins in defining cytoskeletal organization rather than a neuron-specific function in synapse assembly. Septins regulate microtubule organization by preventing catastrophe events, guiding bundling of perinuclear microtubules, and ensuring proper targeting of peripheral microtubules^65^.

### Transcriptomic Adaptations

RNA-seq analyses link the observed structural phenotypes to coordinated transcriptional changes. Microtubule-stabilizing factors, including tau and ringer, are strongly upregulated, whereas γ-TuRC components, kinesin motors, and tubulin-modifying enzymes (TTLL family, Nna1) are downregulated. Actin regulators and extracellular remodeling enzymes are differentially expressed. Notably, core tubulin acetylation regulators HDAC6 and αTAT1 remain unchanged, indicating that increased tubulin acetylation arises from enhanced microtubule stability rather than transcriptional modulation of the acetylation machinery itself.

These transcriptional signatures are consistent with a model in which septin complexes normally constrain microtubule stabilization and preserve cytoskeletal heterogeneity. Upon loss of Sep2 and Sep5, cells shift toward a stability-dominant cytoskeletal state: microtubules persist while nucleation and motor-driven redistribution are suppressed, and spatial microtubule regulation is reduced. Upregulated tau may further prolong polymer lifetime, allowing αTAT1 to acetylate long-lived polymers.

However, these relationships are currently correlative. The extent to which individual factors - particularly tau and ringer-are necessary or sufficient to drive microtubule hyperstabilization in septin mutants remains to be determined through genetic epistasis and functional rescue experiments. Thus, the transcriptomic profile likely represents a combination of primary and compensatory responses to septin complex disruption.

### Functional Consequences: Behavior and Rescue

At the organismal level, septin double mutants exhibit impaired larval locomotion: double mutants move slowly, coil, and display abnormal body orientation. Given the presynaptic defects in vesicle trafficking and active zone organization, these behavioral abnormalities are most consistent with compromised neurotransmission. Similar phenotypes are reported in septin-null nematodes^67^highlighting a conserved role of septins in neuronal development and motor behavior. However, contributions from muscle microtubule disorganization or nuclear positioning defects cannot be excluded, underscoring the multi-tissue impact of septin loss.

Expression of Sep2::GFP partially rescues microtubule organization, vesicle recycling, and locomotion, confirming a direct role for septins in coordinating cytoskeletal integrity and motor output. The incomplete rescue of nuclear positioning suggests tissue-specific sensitivity or dosage-dependent effects of septin composition. Furthermore, allele-specific perturbations reveal that misincorporation versus complete loss of septin subunits disrupts NMJ structure and microtubule organization via distinct but convergent mechanisms.

### Integrated Model and Conclusion

This study utilizes Drosophila genetics to dissect paralog-specific septin functions, revealing redundancy, combinatorial roles, and cell-autonomous effects. Integration of NMJ, muscle, epithelial, behavioral, and transcriptomic analyses enables direct linkage of molecular perturbations to phenotypic outcomes, thereby strengthening mechanistic inference. The conservation of transcriptional and structural effects across neuronal and non-neuronal tissues underscores the broad relevance of septin-mediated cytoskeletal regulation. However, differences in septin isoform complexity and compensatory mechanisms in mammals may limit direct extrapolation to human disease. In addition, the causal contributions of upregulated tau/ringer or downregulated γ-TuRC components remain to be tested genetically. Direct visualization of microtubule dynamics by live imaging will be essential to further refine mechanistic resolution.

Collectively, these findings demonstrate that Sep2 and Sep5 coordinate the microtubule lattice, presynaptic architecture, vesicle trafficking, and muscle microtubule organization. In the absence of Sep2 and Sep5, cells shift toward a rigidity-biased, hyperstabilized state while maintaining overall synaptic architecture. Septin complexes, therefore, maintain a balanced distribution of stable and dynamic microtubule subsets, ensuring functional output without precipitating structural collapse. Elevated tau may further reinforce polymer persistence, facilitating αTAT1-mediated acetylation of long-lived microtubules and consolidating the stability-dominant state.

In conclusion, these findings establish a mechanistic framework in which septin complex composition may specify subsets of microtubules, preserve cytoskeletal heterogeneity, and sustain neuronal structural homeostasis. By constraining excessive stabilization and maintaining balanced microtubule dynamics, septin complexes may buffer cells against transitions toward rigidity-dominant cytoskeletal states that impair membrane trafficking and synaptic function. Together, our results define combinatorial, paralog-specific, and tissue-wide roles for septins as regulators of microtubule state and coordinating cytoskeletal integrity *in vivo*.

## Materials and Methods

### Fly stocks and husbandry

*Drosophila* stocks were maintained on standard fly food under a 12-hour light/dark cycle at 25°C. The following stocks were obtained from the Bloomington *Drosophila* Stock Center: *w*;;;* (BL5905)*, w*;; Sep2¹/TM6B* (BL91002), *w*;; Sep2²/TM6B;* (BL91003), *w*; Sep5²;;* (BL91001). A double mutant stock, (*w*; Sep5²/CyO[tb]; Sep2²/TM6B;)* was generated by crossing single mutants in this study. For *Sep2* expression pattern analysis, the endogenously tagged line *y^1^w*;; P{w^+mC^=Sep2.GFP.SG}3;* (BL26257) was used. Age-matched *y^1^w*;;;* flies were used as genetic background controls. For Sep5 expression analysis, *w*; Sp/CyO; P{w^+mC^=UASp-Sep5.GFP}3/TM6B* (BL51344) flies were crossed to the pan-neuronal driver *y^1^w*;; P{w^+m*^=nSyb-GAL4.S}3/TM6B* (BL51635). Progeny carrying both *UAS-Sep5-GFP* and *nSyb-GAL4* were used to analyze neuronal expression patterns.

To investigate the roles of *Sep2* and *Sep5* in synaptic development, we utilized previously generated null alleles for both genes^3^. The *Sep5²* allele contains a 1558 bp deletion in the Sep5 gene, while the Sep2² allele carries a 1640 bp deletion in the *Sep2* gene. We established a stable *w*^1118^*; Sep5²/CyO[tb]; Sep2²/TM6B;* double mutant stock to rule out the possibility of Sep5 compensating for Sep2. The newly generated line, hereafter referred to as the double mutants, facilitated the selection of double-homozygous mutants at the larval stage. The generation of double mutants was based on the hypothesis that it would adversely affect the overall septin filament, thus it would be unable to function properly. For the generation of double mutants resulting in the loss of function of septins, we selected the largest deletions available in both genes. Another *Sep2* mutant allele that has been used in this study, *Sep2¹*, harbors a 1087 bp deletion in *Sep2* gene. Although the size of the deletion was smaller, flies homozygous for the *Sep2¹* deletion showed markedly reduced viability, with few surviving to adulthood compared to those with the *Sep2²* allele. Consequently, creating double mutants with *Sep5* and *Sep2¹* was not feasible.

For rescue experiments, the endogenously tagged *Sep2* line *y^1^w*;; P{w^+mC^=Sep2.GFP.SG}3* (BL26257) was recombined onto the *Sep2²* mutant chromosome *w*;; Sep2²/TM6B;* (BL91003). The resulting recombinant stock, *w*;; P{w[+mC]=Sep2-GFP.SG}3, Sep2²/TM6B\**, was subsequently used to generate a double mutant rescue line. To this end, the recombinant *Sep2-GFP, Sep2²* stock was crossed into the *Sep5²* background, yielding the rescue genotype *w*; Sep5²/CyO[tb]; P{w[+mC]=Sep2-GFP.SG}3, Sep2²/TM6B;*. In this line, *Sep2* is reintroduced as a GFP-tagged allele under its endogenous regulatory elements, thereby mimicking native *Sep2* expression levels and spatial distribution in an otherwise *w*; Sep5²/CyO[tb]; Sep2²/TM6B;* double mutant background.

### Immunostaining of larval and adult brains and salivary glands

Adult and larval brains and salivary glands were dissected in a PBS buffer, fixed in cold 4% paraformaldehyde (PFA) for 20 minutes at room temperature, and washed three times with PBS containing 0.05% Triton X-100 for 20 minutes. Samples were then blocked in 5% normal goat serum (NGS) in PBS with 0.3% Triton X-100. Subsequently, samples were incubated overnight at 4°C with the primary antibodies in 5% NGS in PBS, 0.3% Triton X-100. After incubation, samples were washed three times with PBST (PBS with 0.05% Triton X-100) for 20 minutes. Secondary antibodies were applied overnight at 4°C. Brain samples were mounted using Vectashield Mounting Medium (Vector Laboratories, Burlingame, CA, USA). For adult brain samples, the bridge mounting method was used.

### Immunostaining of larval fillet

Third instar larvae were dissected in HL3 buffer, fixed in cold 96% methanol, washed three times with PBS containing 0.05% Triton X-100 for 20 minutes, and then blocked in 5% NGS in PBS with 0.3% Triton X-100. Larval fillets were incubated overnight in primary antibodies at 4°C in 5% NGS in PBS, 0.3% Triton X-100. After incubation, samples were washed three times with PBST (0.05% Triton X-100 in PBS) for 20 minutes, followed by a 4-hour incubation in PBST (0.05% Triton X-100 in PBS) at 4°C. Secondary antibodies were applied at 4°C overnight. Following washing three times with PBST (0.05% Triton X-100 in PBS) for 20 minutes, and overnight incubation in PBST (0.05% Triton X-100 in PBS) at 4°C, larval fillets were mounted using Vectashield Mounting Medium (Vector Laboratories, Burlingame, CA, USA).

### Microtubule profiling

Fluorescence intensity distributions of DAPI and tubulin (α-tubulin or acetylated tubulin) were quantified from confocal images using the Radial Profile Plot plugin (Baggethun, 2009) in Fiji. For each nucleus, the nuclear center was defined as the origin (0 µm), and fluorescence intensity was measured along radial profiles extending 100 pixels for each genotype. Distances were converted and expressed in micrometers (µm). For each genotype and imaging channel, multiple radial profiles were collected and exported as tabular data. All downstream analyses were performed in R (version ≥ 4.2) using the *dplyr*, *tidyr*, *ggplot2*, and *scales* packages. At each distance point, replicate intensity values were pooled and summarized as mean fluorescence intensity ± standard error of the mean (SEM). To visualize spatial distributions relative to the nuclear center, intensity profiles were mirrored across the origin, such that for each distance *x*, a corresponding value at –*x* was generated.

Three-dimensional representations of fluorescence intensity distributions were additionally generated using the Interactive 3D Surface Plot plugin for Fiji (Barthel, 2015; version 2.4). Surface plots were created from maximum-intensity projection images using default rendering parameters, with a grid size of 1024, smoothing of 8.0, a perspective of 0, and lighting of 0.44. These 3D surface plots were used exclusively for visualization and were not included in quantitative analyses.

For genotype comparisons, area-under-the-curve (AUC) values were calculated from individual radial intensity profiles. Statistical comparisons between genotypes were performed using the Wilcoxon rank-sum test.

### Nuclear positioning

Myonuclear spacing and muscle area were quantified using a custom Python script implemented with OpenCV (cv2), NumPy, and standard mathematical libraries. Analyses were performed on color-encoded 2D images in which nuclei and muscle boundaries were pre-labeled using distinct pixel colors. The muscle area was estimated by defining a quadrilateral region based on detected boundary pixels. Four representative boundary points were selected from the border pixel set and used as vertices of the polygon. The enclosed area was calculated using the polygon (shoelace) formula.

To quantify nuclear spacing, nuclear coordinates were sorted, and Euclidean distances were calculated between consecutive nuclei to generate a distribution of inter-nuclear “neighbor” distances. From this distribution, mean, minimum, and maximum distance values were extracted. Distances were normalized to muscle size by dividing by the square root of muscle area. For quality control, the script generated annotated images overlaying the muscle boundary, nuclear centers, and lines connecting neighboring nuclei, with the shortest and longest inter-nuclear distances highlighted.

### FM1-43 labeling

Open fillet dissections of wandering third instar larvae were performed in Ca^2+^-free HL3.1 as described earlier^69^ . After dissection, samples were washed twice with HL3 buffer for two minutes each to remove residual tissue debris. FM1-43 dye was prepared at a working concentration of 4 µM and applied in HL3 buffer supplemented with 90 mM KCl. Samples were incubated in the dark for 1 minute, followed by 5 washes with HL3 buffer (2 minutes per wash). After washing, samples were immediately imaged.

### Dyes

DAPI (1 µg/ml; Sigma-Aldrich) was used to label nuclei, and Alexa Fluor™ Plus 555 Phalloidin (Invitrogen, Thermo Fisher Scientific; Cat# A30106) was used to label actin filaments. FM1-43 dye (Invitrogen, Carlsbad, CA, USA; T-35356 or T-3163) was used for synaptic vesicle retrieval.

### Primary antibodies

Following primary antibodies have been used: Mouse Acetyl-tubulin (66200-1-Ig, ProteinTech) (1:1000 dilution), Mouse anti-E-cadherin (extracellular domain) (AB_528120, Developmental Studies Hybridoma Bank, University of Iowa (DSHB)) (1:100 dilution), rat 7E8A10 anti-Elav (AB_528218, DSHB) (1:50 dilution), Rabbit anti-GFP (TP401, Torrey Pines)(1:1000), 4A1 mouse anti-Tubulin alpha (AB_2732839, DSHB) (1:200 dilution), Rabbit anti-HRP (Jackson ImmunoResearch Labs Cat# 323-005-021, RRID: AB_2314648) (1:2000 dilution). Rabbit anti-HRP (Jackson ImmunoResearch Labs Cat# 323-005-021, RRID: AB_2314648) (1:2000 dilution), 4F3 mouse anti-DLG1 (AB_528203, DSHB) (1:50 dilution), Mouse anti-Syn (AB_528479, DSHB) (1:25 dilution), Mouse anti-BRP (AB_2314866, DSHB) (1:100 dilution), Mouse anti-Futsch (AB_528403, DSHB) (1:50 dilution), Mouse anti-GluRAII (AB_528269, DSHB) (1:50 dilution). 4C9H4 anti-peanut (AB_528429, DSHB) (1:20 dilution).

### Secondary antibodies

Goat anti-Mouse IgG (H + L), Alexa Fluor® 647 conjugate (Invitrogen, #A-21236) (1:800 dilution), Goat anti-Rat IgG (H + L), Alexa Fluor® 555 conjugate (Invitrogen, #A-21434) (1:800 dilution). Goat anti-Rabbit IgG (H + L), Alexa Fluor® 488 conjugate (Invitrogen, #A-11008 (1:800 dilution)).

### Larval locomotion assay

All flies were maintained at 25°C with a 12-hour light/dark cycle and 45% humidity. Crosses involved 10 males and 10 females to minimize density effects, and adults were transferred to new vials after 5 days. Larvae were collected 5–6 days after egg-laying, sexed, and used for behavioral tests performed separately for males and females.

Fresh 0.8% agar plates made with food agar and distilled water were used. Larvae were washed twice with autoclaved water to remove food residues, sexed, and separated. All behavioral experiments for males and females were conducted independently to investigate sexual dimorphism in larval locomotion, with larvae habituated on agar plates. The genotype order was randomized to reduce bias and account for variations in habituation timing.

Larvae were transferred with a paintbrush to fresh agar plates for locomotion analysis. After confirming active crawling, a 2-minute video was recorded as before. The agar plates were positioned on a light source in a dark room to improve contrast, then cleaned and moistened with distilled water for future tests, avoiding excess water or food residues to ensure natural behavior. For recording, a simple setup involved an iPad roughly 31 cm above the surface. All recordings were made under stable environmental conditions.

Larval locomotion was recorded at room temperature using a custom setup. Fifteen to seventeen larvae were gently placed at the center of 11.5 × 18 cm agar plates with 0.8% agar, 2 mm thick. 2-minute videos were recorded at 30 fps and 1080 × 1920 resolution on a 9th-generation iPad, with backlighting in a dark room. The agar surface was cleaned and moistened between recordings.

Processed videos in Fiji involved adjusting brightness and contrast, inverting colors, and reducing the frame rate to 10 fps. Trajectories were extracted with FIMTrack, with manual corrections for larval collisions. Fiji is used to convert pixels to millimeters for accurate analysis of distance and velocity. Accumulated distance and velocity values are directly obtained from the FIMtrack output. Additional features, such as go-phase percentage, body orientation, and coiling behavior, were calculated using custom Python scripts in PyCharm. Statistical analysis was conducted with GraphPad Prism (v8.4.2), applying the Shapiro–Wilk test for normality and appropriate parametric or nonparametric tests.

Preliminary experiments optimized behavioral assay conditions and verified reproducibility over days. Environmental factors like lighting and surface tilt were tested for stability by observing larval bending, which showed no significant effects. Sexual dimorphism in available larvae was also assessed to account for gender-specific behavioral differences.

Videos were manually edited and analyzed with FIMTrack. Locomotion parameters, including some unspecified metrics, were calculated using a custom Python script available upon request.

Videos were processed in Fiji through brightness/contrast adjustments and color inversion to generate high-contrast sequences at 10 fps, with a black background and white larvae. FIMTrack software extracted locomotion data, with manual identity corrections for collided larvae.

Distance and velocity data from FIMTrack were processed manually. The final pixel-based distance and mean velocity were recorded over 1200 frames. Additional parameters like go-phase percentage, posture, and body-bending angles were analyzed using a Python script available upon request. Pixel measurements were converted to millimeters and millimeters per second through Fiji calibration. Data visualization and statistical analysis were performed in GraphPad Prism (v8.4.2).

### Light microscopy and tissue melanization

Salivary glands were examined under a light microscope (Leica) at the highest magnification. Images were captured using hand-held afocal photography, with an iPhone 15 (Apple Inc., Cupertino, CA, USA) camera positioned directly against the microscope eyepiece.

### Quantification of bouton numbers

The number of boutons was counted using the Cell Counter plugin in ImageJ 1.00.

### Quantification of the spatial distribution of pre- and post-synaptic markers

To quantify the NMJ phenotype, we used an in-house protocol and Python script to measure the area covered by each protein relative to the whole presynaptic area.

Images were processed in ImageJ 1.00 by applying binarization, thresholding, and despeckling to enhance signal clarity. Signals from neighboring NMJs were manually removed to ensure accurate quantification. The processed channels containing only positive signals for the respective antibodies were then merged, and colocalization was quantified using the Color Counter plugin.

The spatial distribution percentage of synaptic proteins over the synaptic area was calculated using the following formula and in house generated Python script based on the following formula: Number of double-positive pixels for the target synaptic protein and HRP/Total number of HRP-positive pixels ×100. Total number of HRP-positive pixels/Number of double-positive pixels for the target synaptic protein and HRP ×100. For postsynaptic protein DLG1, normalization was performed relative to the muscle area.

### Quantification of FM1-43 dye uptake by the presynaptic membrane

FM1-43 dye uptake images were processed in 8-bit mode in ImageJ 1.00. The bouton area and a random background area were selected, and the average pixel intensities (ranging from 0 to 255) were calculated. The fold change in dye uptake was determined using the following formula and using an in-house Python script based on the following formula: Average pixel intensity in the bouton area / Average pixel intensity in an equivalent-sized muscle area not covered by the NMJ.

### Calculation of the distance between myonuclei and determining the shape of nuclei within the body-wall muscles of larvae

To evaluate nuclear architecture, larval muscles were stained with DAPI, and confocal images were acquired. Morphometric analyses were conducted using Fiji software and a custom Python script.

To quantitatively assess changes in nuclear spacing, we used nearest-neighbor analysis. This method calculates the shortest distance from the center of each nucleus to its nearest neighbor, providing an average measure of inter-nuclear proximity.

### Imaging

Larval and adult brains and NMJs were imaged using a Leica TCS SP5 confocal microscope (Leica Microsystems GmbH, Wetzlar, Germany). Identical imaging settings were used for all samples, with z-stack images (1.5 µm step size) and 1024 × 1024-pixel resolution. FM1-43 staining was imaged using System I, with 514 nm - Alexa 633 excitation/emission parameters.

### RNA extraction and library preparation

Total RNA was extracted from four technical replicates per genotype. Total RNA was isolated from *Drosophila melanogaster* 12-15 male larvae. Male larvae were collected on a petri dish under a dissection microscope and transferred into an Eppendorf tube. Larvae were snap-frozen in liquid nitrogen and immediately homogenized using RNase-free pellet pestles. 400 μL TRIzol reagent (Thermo Fisher Scientific) was added, and homogenization was repeated. An additional 400 μL of TRIzol was then added, and the lysate was sheared by passing it through a 20-gauge needle fitted to a syringe at least 20 times. After a 5-minute incubation at room temperature, 160 μL of chloroform was added, the mixture was vortexed vigorously for 15 seconds, and then it was incubated for an additional 3 minutes. Phase separation was performed after centrifugation at 13,200 × g for 15 minutes at 4°C. The aqueous (upper) phase was carefully transferred to a new RNase-free tube. RNA was precipitated by adding 400 μL of cold isopropanol, incubating at room temperature for 10 minutes, and then centrifuging at 13,200 × g for 15 minutes at 4°C. The pellet was washed once with 1 mL of 75% ethanol, centrifuged for 5 minutes, air-dried, and resuspended in 20 μL of RNase-free water. Samples were incubated at 55°C for 10 minutes to aid resuspension and stored at −80°C. Finally, Nanodrop quantification was performed, and standard purity criteria (A260/A280 ≈ 1.8–2.0 and A260/A230 ≥ 2.0) were applied to ensure sample quality. Enrichment for poly-A mRNAs and subsequent library construction were carried out using the Watchmaker mRNA Library Prep Kit in accordance with the manufacturer’s guidelines.

### RNA-sequencing and data analysis

Raw sequence reads were processed and mapped to the *Drosophila melanogaster* reference genome (dm6) using the STAR package. Count matrices were analyzed using the edgeR (v3.42.0) Bioconductor package in R (v4.3.1). To account for compositional differences between libraries, normalization factors were calculated using the Trimmed Mean of M-values (TMM) method. Genes with low expression counts across samples were filtered using the filterByExpr function.

A generalized linear model (GLM) was fitted to the data to estimate common, trended, and tagwise dispersions. Differential expression (DE) was determined using the likelihood ratio test (LRT). P-values were adjusted for multiple testing using the Benjamini-Hochberg (BH) false discovery rate (FDR) procedure. Genes were defined as significantly differentially expressed (DEGs) if they exhibited an FDR < 0.05 and an absolute log2 fold-change (|log2FC|) > 1.0.

Gene Ontology (GO) enrichment analysis was performed using the clusterProfiler package. To investigate the specific impact on the cellular architecture, we performed a targeted analysis of cytoskeletal and organization networks. Gene lists were derived from the GO:0005856 (cytoskeleton) and GO:0007010 (cytoskeleton organization) hierarchies using the org.Dm.eg.db annotation package.

Heatmaps were generated using the pheatmap package, applying row-wise Z-score normalization. Volcano plots were generated using ggplot2 and ggrepel to visualize the distribution of fold changes and statistical significance across all contrasts.

To characterize the global and pathway-specific transcriptomic trajectories of the single and double mutants, Principal Component Analysis (PCA) was performed using the prcomp function in R. PCA was applied to the log-transformed Counts Per Million (log-CPM) expression matrix following TMM normalization.

To ensure the biological signal was not confounded by genomic noise or subset size bias, two distinct PCA frameworks were utilized: To characterize the primary axes of transcriptomic divergence and verify the consistency of biological replicates, we performed Principal Component Analysis (PCA) using the top 500 genes with the highest row-wise variance across all samples. This global approach enabled us to assess the overall landscape of gene expression changes and identify the major drivers of variation between wild-type and mutant genotypes. To further isolate the variance specifically attributable to the cytoskeletal regulatory network, a Targeted Cytoskeleton PCA was conducted using all detected genes associated with the Gene Ontology (GO) terms for Cytoskeleton (GO:0005856) and Cytoskeleton Organization (GO:0007010).

### *In silico* modelling of mutant Sep2¹ protein and Sep2² proteins

The protein structure of *Drosophila melanogaster* Sep2RA (UniProt ID: P54359; SEPT2_DROME), the canonical amino acid sequence was retrieved directly from UniProt, and was predicted using AlphaFold2 (AF2). For the alternative isoform, Sep2RB, and two putative mutant variants (Sep2^1^ and Sep2^2^), genomic DNA sequences were obtained from FlyBase^70^. The longest open reading frames (ORFs) were identified using the ORF Finder tool (https://www.bioinformatics.org/sms2/orf_find.html), and the resulting predicted protein sequences were translated accordingly. For Sep2^1^ and Sep2^2^, deletions were derived from their original publication^29^. All input sequences were provided to the AF2. AF2 models were run via ColabFold using MMseqs2^71–73^, selecting the mmseqs2_uniref_env MSA mode and the unpaired_paired pairing mode. The alphafold2_ptm model was used, with the number of recycles adjusted to 6. AF2 predictions were evaluated using the confidence metrics predicted Local Distance Difference Test (pLDDT) and Predicted Aligned Error (PAE).

### *Drosophila melanogaster* Sep1-Sep2^1^ complex structure

The crystal structure of the Sep1–Sep2 complex (PDB ID: 8DKT)^45^. The only available experimental septin structure of *Drosophila melanogaster* in the PDB was extracted. In the experimentally resolved 8DKT complex, chain A (Sep1) spans residues 34–304, and chain B (Sep2) spans residues 41–308. The internal missing loop regions (residues 215–217, 244–251 in chain A and residues 90–95, 248–251 in chain B) were modelled with the MODELLER^74^ plugin in ChimeraX^75,76^, generating ten loop models and choosing the model with the lowest zDOPE score passing geometry/validation^77^. The Sep2^1^ putative mutant protein, encoding residues 171–419, was generated by truncating Sep2 (chain B) of the 8DKT structure at residue 171 to construct the Sep1–Sep2^1^ complex. GDP bound to Sep1 in the 8DKT structure was retained in the Sep1–Sep2^1^ complex, whereas GTP and Mg²⁺ associated with Sep2 were not included, as ligand binding by Sep2^1^ is suspicious. The XRC-truncated Sep2^1^ structure was further validated against predictions for the AF2 monomer^71^.

### Molecular dynamics system preparation

All-atom molecular dynamics (MD) simulation was performed to analyze the constructed *Drosophila melanogaster* Sep1–Sep2^1^ complex. Crystal waters and buffer contaminants were removed from the complex structure, while the Sep1-bound GDP was retained. The Sep1-Sep2^1^ complex was solvated in an orthorhombic water box with edge distances 111.0 Å, ensuring a minimum solute–box distance of 10 Å in all directions. To neutralize the system and approximate physiological ionic strength, 123 Na⁺ and 115 Cl⁻ ions were introduced to reach a final salt concentration of 150 mM. The system contained 1.282 × 10⁵ atoms. System setup and solvation/ionization were performed using the Solvation Builder module in CHARMM-GUI^78^. All interactions were treated using the CHARMM36m force field^79–82^. Production simulations were performed using the NAMD 3.0 engine^81–83^. The TIP3P water model was used to explicitly treat the solvent^84^. The system was simulated under three-dimensional periodic boundary conditions with a 2-fs integration timestep. The particle-mesh Ewald (PME) method was used to compute long-range electrostatics (grid spacing of 1 Å), and real-space nonbonded interactions were truncated at a 12 Å cutoff^85^. The system energy was minimized for 10.000 steps, then equilibrated in the NVT ensemble at 310 K for 250 ps. An over-600-ns production trajectory was obtained under NPT conditions at 310 K and 1 atm, with temperature control provided by a Langevin thermostat and pressure regulation by a Langevin piston^86–88^.

### Trajectory analysis

The production trajectory of the Sep1–Sep2^1^ complex was analyzed using pairwise root-mean-square deviations (RMSD) and root-mean-square fluctuations (RMSF) over Cα atoms to assess global structural changes and residue-level flexibility, respectively. Analyses were performed with in-house Python scripts using the MDAnalysis library^89^. Interface SASA (iSASA) was computed with in-house VMD scripts by subtracting the SASA of the Sep1-Sep2^1^ complex from the sum of the SASAs of the isolated Sep1 and Sep2^1^ subunits. Principal component analysis (PCA) was performed using the Bio3D package in R^90,91^. Structures and trajectories were visualized using ChimeraX^75,76^ and VMD^92^.

## Acknowledgement

The monoclonal antibodies DE-Cadherin (T. Uemura, Kyoto University), Elav (G.M. Rubin, Janelia Farm/HHMI), α-Tubulin (M.T. Fuller, Stanford University), Discs large and DGluR-IIA (C. Goodman, University of California, Berkeley), Synapsin and Bruchpilot (E. Buchner, Universitätsklinikum Würzburg), Futsch (S. Benzer and N. Colley, California Institute of Technology), and Peanut (G.M. Rubin, University of California, Berkeley) were obtained from the Developmental Studies Hybridoma Bank, created by the NICHD of the NIH and maintained at The University of Iowa, Department of Biology, Iowa City, IA 52242. Stocks obtained from the Bloomington *Drosophila* Stock Center (NIH P40OD018537) were used in this study.

The authors would like to thank Gülay Kaya, Aleyna Karbuz, Pınar Aydan Kekeç, and Mustafa Kaan Kozluca for their assistance with immunostaining experiments.

## Author contributions

F.L., T.A., and A.C. designed research; F.L., T.A., A. S., I.S., R.E., and G.K.P. performed research; F.L., T.A., A.S., I.S., R.E., and G.K.P. analyzed data; F.L., T.A., A.S., I.S., G.K.P., and A.C. wrote the paper.

---

**Supplementary Figure 1.**
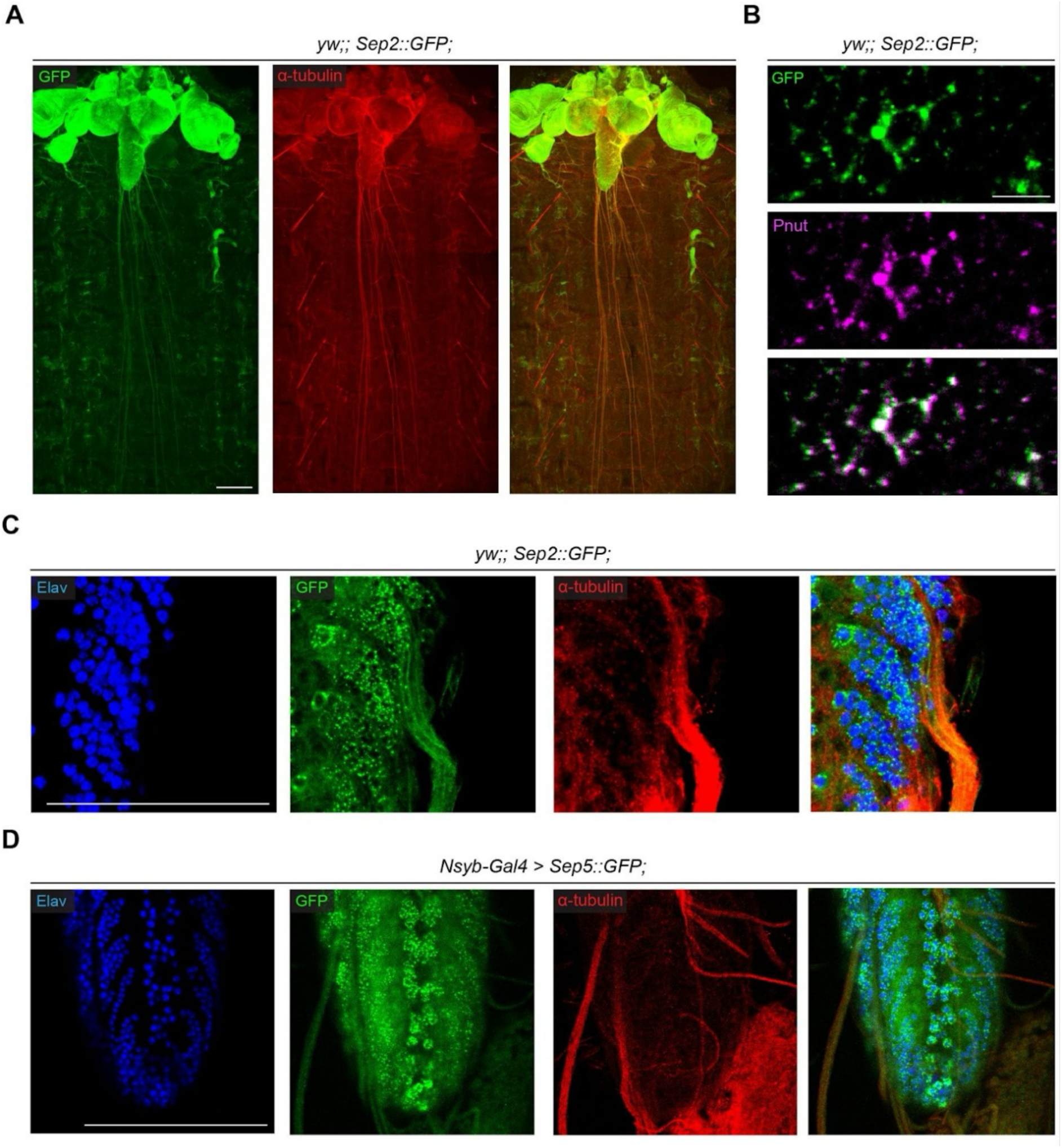
Sep2 and Sep5 localize to neuronal cell bodies, axons, and distal axonal branches in Drosophila melanogaster. (A) Confocal images of a third-instar larval fillet showing the ventral nerve cord (VNC) and associated axonal tracts from a *yw;; Sep2::GFP;* larva, in which the endogenous sep2 coding sequence is tagged with EGFP under its native promoter. Sep2::GFP (green) is enriched along axonal bundles and neuronal regions marked by α-tubulin (red), showing overlapping signals along axons. Scale bar: 20 µm. (B) Adult brain from *yw;; Sep2::GFP* flies stained for Pnut (magenta), a core septin complex component. Sep2::GFP and Pnut display overlapping punctate patterns within neuronal cell bodies. Scale bar: 5 µm. (C) Maximum-intensity projection of the larval ventral nerve cord stained with DAPI (blue; nuclei), Sep2::GFP (green), and α-tubulin (red). Sep2::GFP shows both punctate cytoplasmic localization and enrichment along tubulin-positive axonal structures within the VNC. Scale bar: 20 µm. (D) Maximum-intensity projection of the larval ventral nerve cord expressing Sep5::GFP under pan-neuronal control (*nSyb-Gal4 > Sep5::GFP*). Sep5::GFP (green) localizes to neuronal cell bodies marked by Elav-positive nuclei (blue) and to axonal regions overlapping with α-tubulin (red). Sep5::GFP displays a punctate pattern similar to that observed for Sep2::GFP. Scale bar: 20 µm.

**Supplementary Figure 2.**
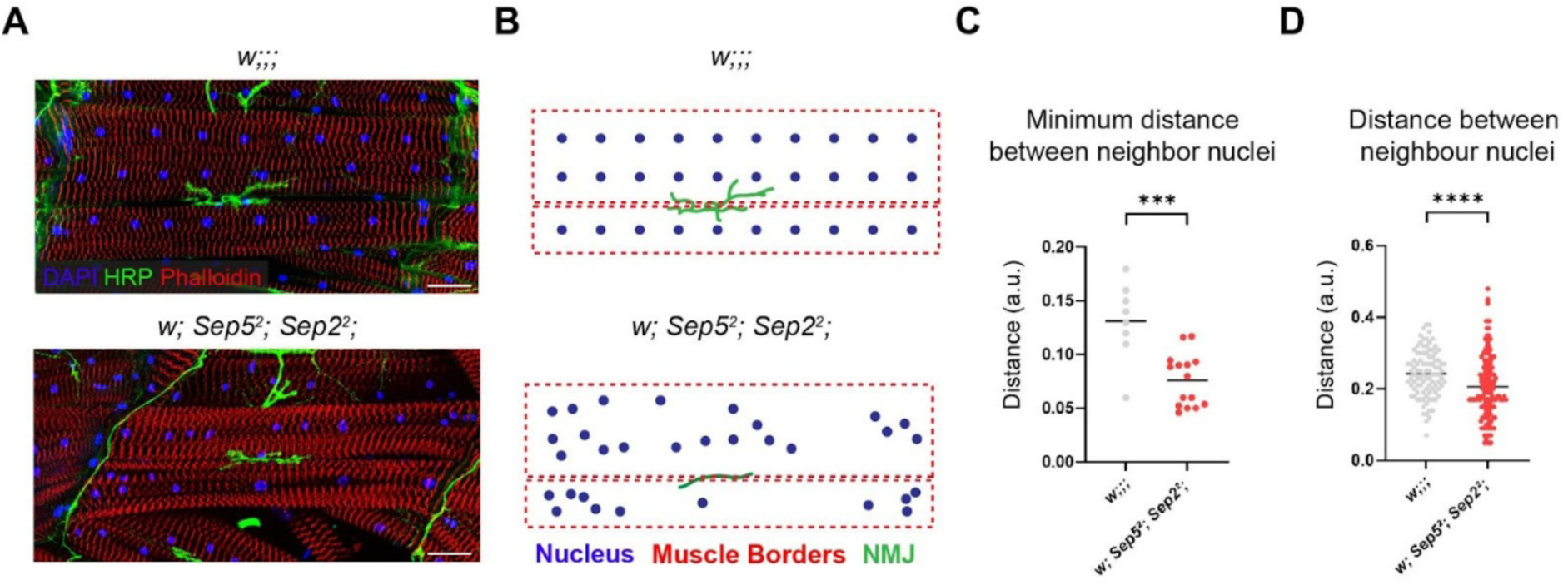
Loss of *Sep2* and *Sep5* disrupts myonuclear spacing in *Drosophila* larval muscles. (A) Representative confocal images of larval muscle 6 (segment A4) stained with DAPI (blue; nuclei), anti-HRP (green; motor neuron membrane), and phalloidin (red; F-actin) in control (*w;;;*) and double mutant (*w; Sep5²; Sep2²*) larvae. While control muscles display evenly spaced myonuclei along the muscle fiber, double mutants exhibit irregular nuclear positioning and local clustering. Scale bar: 40 µm. (B) Schematic representation of myonuclear distribution and neuromuscular junction (NMJ) positioning in control (top) and double mutant (bottom) muscles. Blue dots indicate myonuclei, green structures represent NMJ terminals, and red dashed lines mark muscle borders. Control muscles show regular nuclear spacing, whereas double mutants display disrupted spatial organization with clustered nuclei. This panel is schematic. (C) Quantification of the minimum distance to the nearest neighboring nucleus (nearest-neighbor distance) in muscle 6, normalized to muscle area (a.u.). Each data point represents one larva (n > 7 larvae per genotype). (D) Quantification of distances between neighboring nuclei in muscle 6 normalized to muscle area (a.u.). Each data point represents an individual nucleus pooled from at least 7 larvae per genotype. Data are shown as individual values with mean ± SEM. Statistical analysis: unpaired two-tailed t-test after Shapiro–Wilk normality test; ***p < 0.001, ****p < 0.0001.

**Supp Figure 3.**
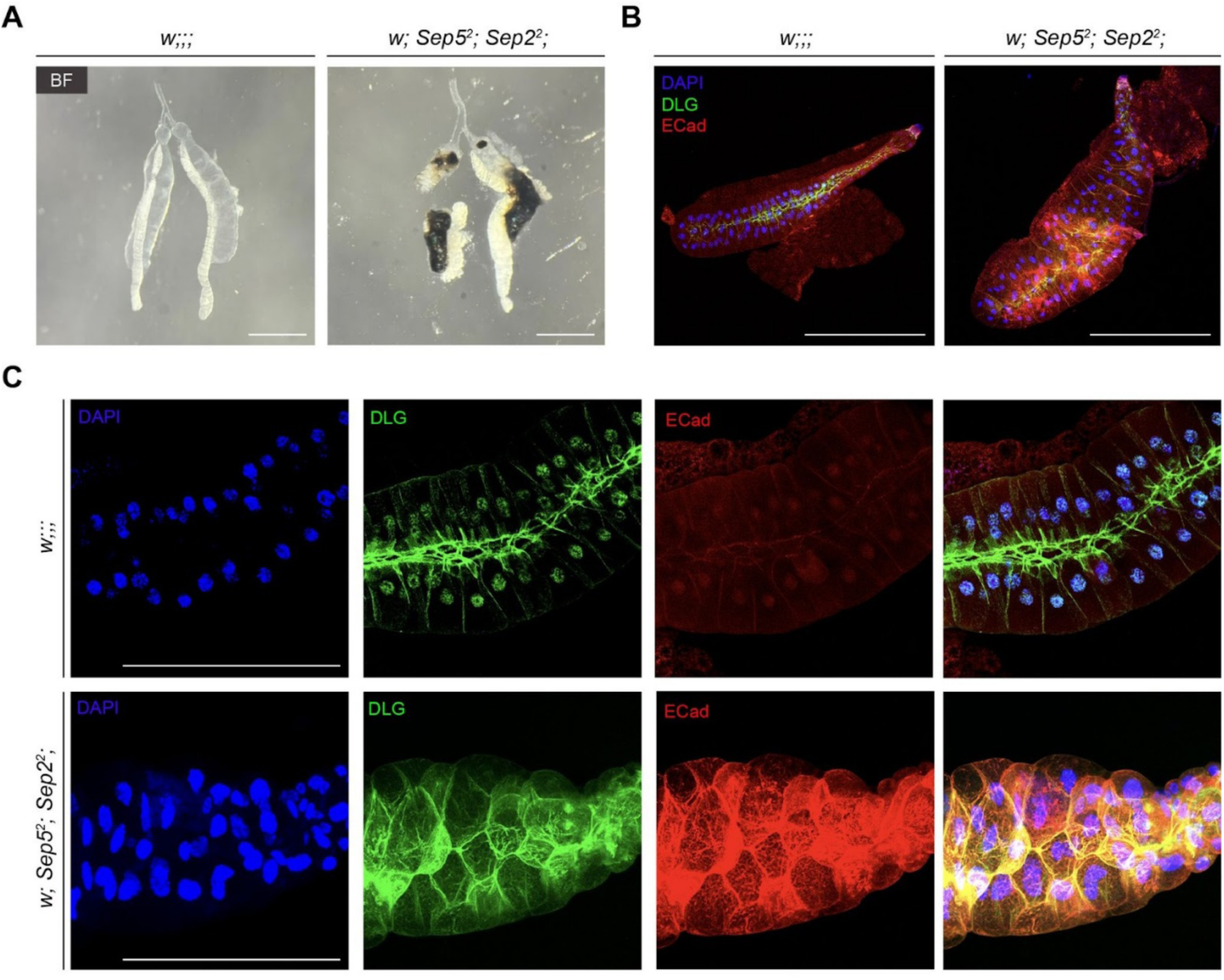
Loss of Sep2 and Sep5 disrupts the morphology and organization of the epithelial cells in the larval salivary gland. (A) Brightfield (BF) images of third instar larval salivary glands from control (*w;;;*) and double mutant (*w; Sep5²; Sep2²*) larvae. Salivary glands of double mutants appear morphologically different, with signs of tissue melanization (black area). (B) Confocal images of salivary glands stained with DAPI (blue; nuclei), anti-E-cadherin (green; E-cad), and anti-DLG1 (red; Discs Large) in control and double mutant larvae. Note the irregular shape and size of the salivary gland and its defective tubular structure, including the disorganized structure of the cell borders and nuclei. (C) High-magnification images of the epithelial cell layer show defects in the localization and distribution of E-cadherin and DLG1 in salivary glands of double mutants. Nuclear positioning, shape, and size are also defective in double mutants. Scale bars: 20 μm (all panels).

**Supplementary Figure 4.**
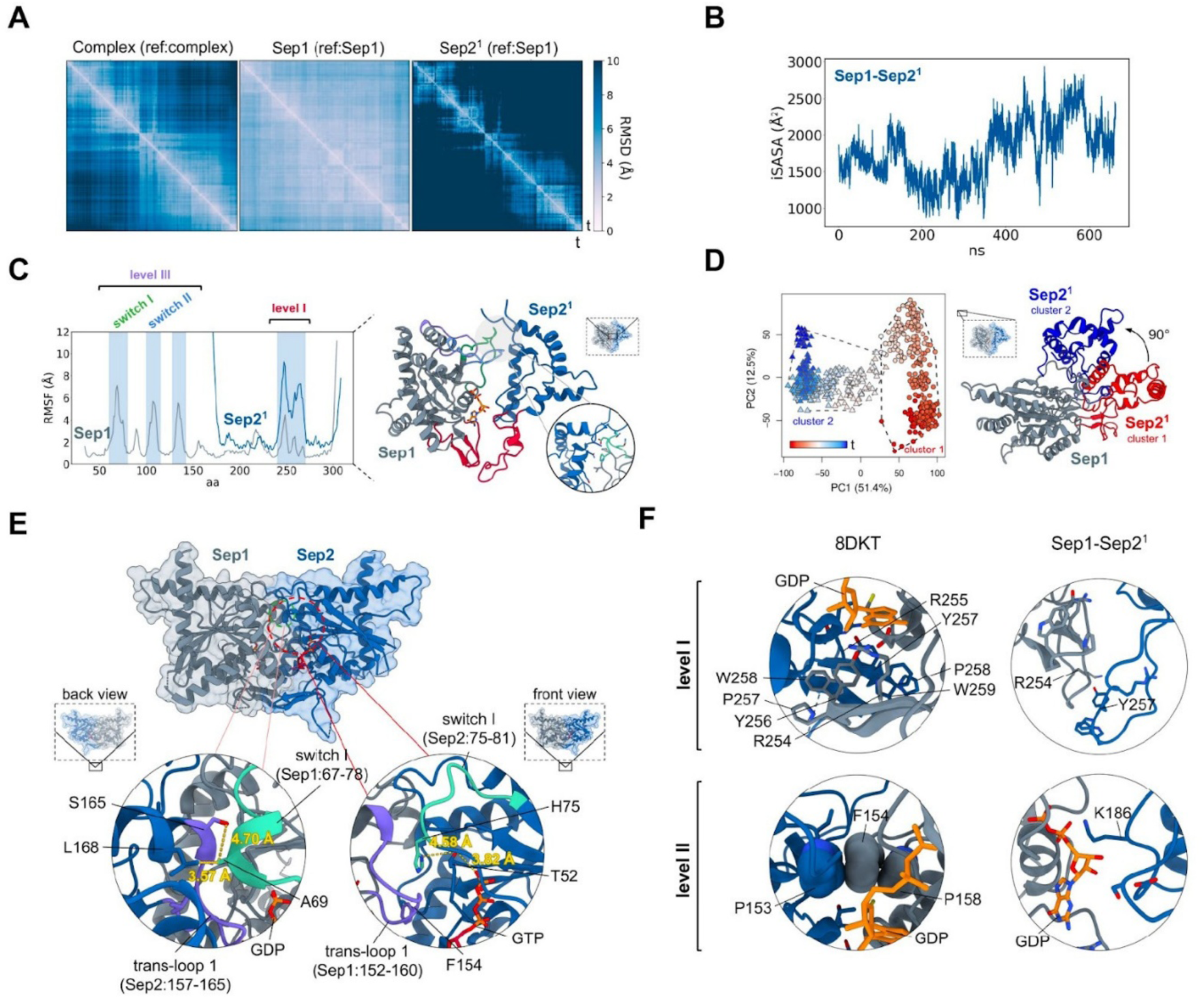
Multi-level Sep1–Sep2 interaction interfaces in the XRC structure. (A) Pairwise time-resolved RMSD matrices (Å) computed on Cα atoms relative to the initial frame. Left: RMSD of the Sep1–Sep2¹ complex after alignment to the full complex structure. Middle: Sep1 RMSD after aligning the complex to Sep1. Right: Sep2¹ RMSD after aligning the complex to Sep1. (B) Interface SASA (iSASA) of the Sep1–Sep2¹ complex, computed by subtracting the complex SASA from the sum of individual Sep1 and Sep2¹ SASAs. Cα RMSF for Sep1 and Sep2¹, with each subunit aligned to itself. Regions with RMSF > 4 Å (Sep1: switch I, switch II (level III) and level I; Sep2¹: level I) are mapped onto the Sep1–Sep2¹ complex representative from the second MD cluster. (D) PCA of the Sep1–Sep2¹ complex showing PC1 and PC2 (explained variance indicated), together with k = 2 clustering. Representative complex structures of the clusters are shown, with time-based coloring from red to blue. (E) Level III interactions in the 8DKT XRC structure, mediated by key motifs (switch regions and trans-loops) that are missing in the Sep1–Sep2^1^ complex, are depicted. (F) Level I and Level II interactions shown for the 8DKT XRC structure (left) and the Sep1–Sep2^1^ complex extracted from cluster 2 (right).

**Supplementary Figure 5.**
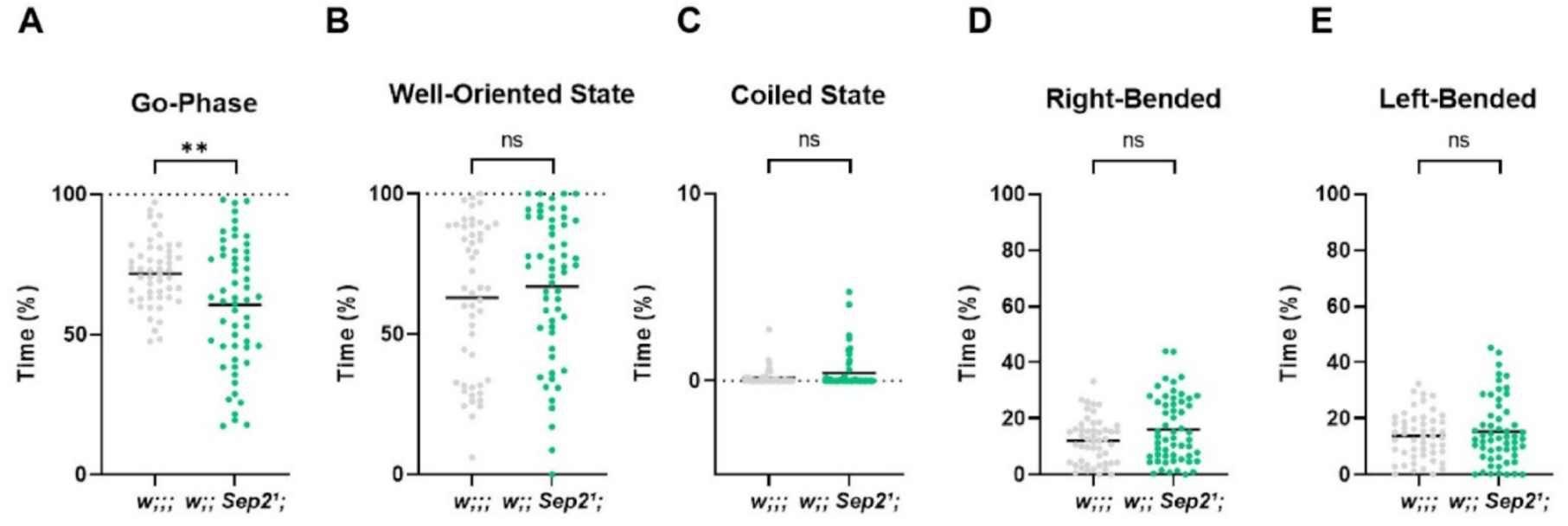
Behavioral state distribution in *Sep2¹* mutants. (A) Quantification of locomotor state occupancy during 120 s recordings of third-instar larvae. The percentage of frames spent in each behavioral state is shown for control (*w;;;*) and *Sep2¹* mutant (*w;; Sep2¹;*) animals. Each dot represents an individual larva, and horizontal bars indicate the mean. (B) *Sep2¹* mutants spend a significantly reduced proportion of time in the go-phase, defined as frames in which larvae actively propel themselves forward. (C) The proportion of time spent in a well-oriented state, defined by proper alignment of anterior, middle, and posterior body segments, is not significantly different between control and *Sep2¹* larvae. (D) Time spent in the coiled state does not differ significantly between genotypes. (E) The percentages of frames exhibiting right-bended (E) or left-bended (F) body postures are comparable between control and *Sep2¹* larvae, indicating no significant bias in lateral bending behavior. Statistical comparisons were performed using Shapiro–Wilk normality testing followed by two-tailed t-tests for parametric data or Mann–Whitney U tests for nonparametric data, as appropriate. **p < 0.01; ns, not significant.

**Supplementary Figure 6.**
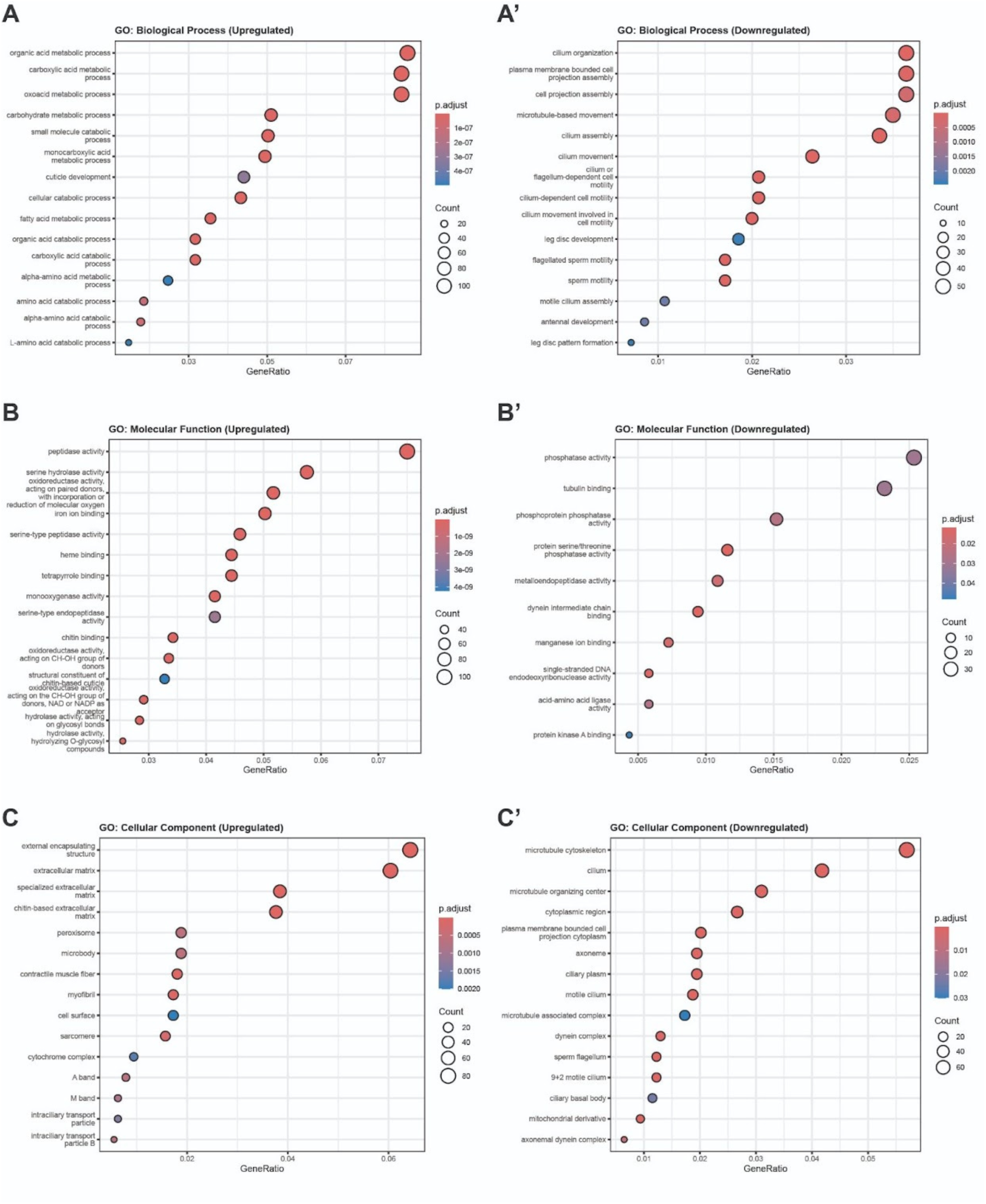
Gene Ontology enrichment analysis of differentially expressed genes. (A–C) Gene Ontology (GO) enrichment analysis of upregulated genes showing significantly enriched Biological Process (A), Molecular Function (B), and Cellular Component (C) categories. Dot plots display the top enriched GO terms ranked by adjusted p-value. The x-axis represents GeneRatio (proportion of input genes annotated to each term), dot size corresponds to the number of genes (Count), and color indicates the adjusted p value (Benjamini–Hochberg correction). Upregulated genes are predominantly enriched in metabolic and catabolic processes (A), including organic acid and carbohydrate metabolism; enzymatic and catalytic activities such as peptidase and oxidoreductase functions (B); and extracellular and structural components, including extracellular matrix and contractile fiber-related terms (C). (A′–C′) GO enrichment analysis of downregulated genes showing enriched Biological Process (A′), Molecular Function (B′), and Cellular Component (C′) categories. Downregulated genes are strongly associated with cilium organization and movement, microtubule-based processes, and cell projection assembly (A′); phosphatase activity, tubulin binding, and dynein-related functions (B′); and microtubule cytoskeleton, ciliary structures, and axonemal components (C′). Adjusted p-values are shown as color gradients; only significantly enriched terms are presented.

**Supplementary Table 1.** Differentially Expressed Genes (adjusted p < 0.01) Enriched in Cytoskeletal Binding and Synaptic Development GO Categories. This table lists significantly differentially expressed genes (DEGs; adjusted p-value < 0.01) identified by RNA-seq analysis and annotated based on Gene Ontology (GO) terms. Genes are grouped according to enriched molecular function categories, actin binding (GO:0003779) and tubulin binding (GO:0015631), and biological process categories including axonogenesis (GO:0007409), endocytosis (GO:0006897), and regulation of neuromuscular junction development (GO:1904396). For each gene, the table provides annotation ID, gene symbol, adjusted p-value, log_2_ fold change, and associated GO functional classification. Together, these data highlight coordinated transcriptional changes affecting cytoskeletal organization, membrane trafficking, and synaptic development pathways.

| Annotation | Symbol | adjusted p-values | log2 fold change | Molecular Function/ biological process |
| --- | --- | --- | --- | --- |
| CG18146 | *NimC2* | 0.00004423429355 | -2.770869025 | Endocytosis |
| CG44533 | *Nna1* | 0.000007257417596 | -2.705065216 | Tubulin binding |
| CG2955 | *CG2955* | 0.000002870190879 | -2.246896549 | Tubulin binding |
| CG7716 | *t-Grip84* | 0.000007072452863 | -2.23205219 | Tubulin binding |
| CG32232 | *t-Grip128* | 0.000009852611411 | -2.191754784 | Tubulin binding |
| CG42840 | *dachs* | 0.000000009152601156 | -2.113608772 | Actin binding |
| CG12983 | *CG12983* | 0.0000005731927936 | -2.108654271 | Tubulin binding |
| CG4952 | *dac* | 0.0000001380327372 | -2.073815013 | Axonogenesis |
| CG1794 | *Mmp2* | 0.0000000005834694272 | -2.05087369 | Axonogenesis |
| CG33287 | *CG33287* | 0.00001218927577 | -2.031397054 | Tubulin binding |
| CG4609 | *fax* | 0.00004452011411 | -1.967635096 | Axonogenesis |
| CG31787 | *CG31787* | 0.00003236652431 | -1.956485588 | Endocytosis |
| CG16716 | *TTLL6A* | 0.0000002452739234 | -1.946633006 | Tubulin binding |
| CG7227 | *CG7227* | 0.005313426066 | -1.934489412 | Endocytosis |
| CG33960 | *Sema2b* | 0.000000007825635172 | -1.910597251 | Axonogenesis |
| CG18109 | *t-Grip91* | 0.00005518878262 | -1.899487819 | Tubulin binding |
| CG10845 | *CG10845* | 0.00002011596244 | -1.850730583 | Tubulin binding |
| CG30177 | *CG30177* | 0.00006421402642 | -1.849908833 | Tubulin binding |
| CG9015 | *en* | 0.000000005843028845 | -1.836543057 | Axonogenesis |
| CG33208 | *Mical* | 0.00001925479478 | -1.801170591 | Actin binding |
| CG33208 | *Mical* | 0.00001925479478 | -1.801170591 | Axonogenesis |
| CG2082 | *CG2082* | 0.000001576401593 | -1.799956041 | Tubulin binding |
| CG31907 | *Mst27D* | 0.0001097627464 | -1.786246761 | Tubulin binding |
| CG10719 | *brat* | 0.00004301538176 | -1.751750802 | Endocytosis |
| CG10719 | *brat* | 0.000043 | -1.75 | NMJ development |
| CG32371 | *CG32371* | 0.0002082646962 | -1.739376258 | Tubulin binding |
| CG32238 | *TTLL1B* | 0.000007607800788 | -1.721385703 | Tubulin binding |
| CG2718 | *Gs1* | 0.0002793617342 | -1.710220629 | Endocytosis |
| CG2577 | *CG2577* | 0.0001708801158 | -1.706795257 | Endocytosis |
| CG6634 | *mid* | 0.000001659212955 | -1.695340269 | Axonogenesis |
| CG1916 | *Wnt2* | 0.000002516299595 | -1.619061838 | Axonogenesis |
| CG42282 | *NimA* | 0.009966654494 | -1.575662635 | Endocytosis |
| CG12147 | *CG12147* | 0.0001137752651 | -1.57451973 | Endocytosis |
| CG9397 | *jing* | 0.00007453596578 | -1.571119001 | Axonogenesis |
| CG17122 | CG17122 | 0.0001364318745 | -1.532361677 | Tubulin binding |
| CG3219 | Klp59C | 0.00003208115965 | -1.525974512 | Tubulin binding |
| CG33286 | CG33286 | 0.001121069667 | -1.522135171 | Tubulin binding |
| CG3166 | aop | 0.0003007436574 | -1.516329884 | Endocytosis |
| CG14271 | Gas8 | 0.0002701122715 | -1.507087588 | Tubulin binding |
| CG8918 | TTLL1A | 0.003977684599 | -1.48827879 | Tubulin binding |
| CG12395 | CG12395 | 0.001204316305 | -1.47991092 | Tubulin binding |
| CG15676 | CG15676 | 0.001373114858 | -1.452108217 | Tubulin binding |
| CG42273 | mnb | 0.0000889 | -1.45 | NMJ development |
| CG42273 | mnb | 0.0000888669369 | -1.448848457 | Endocytosis |
| CG7131 | CG7131 | 0.0001355416098 | -1.444496666 | Tubulin binding |
| CG17838 | Syp | 0.0000292 | -1.44 | NMJ development |
| CG42338 | Ten-a | 0.002951087657 | -1.431325085 | Axonogenesis |
| CG42338 | Ten-a | 0.00295 | -1.43 | NMJ development |
| CG12192 | Klp59D | 0.00007569393377 | -1.404064519 | Tubulin binding |
| CG4889 | wg | 0.0000148 | -1.4 | NMJ development |
| CG7892 | nmo | 0.000165 | -1.4 | NMJ development |
| CG15306 | CG15306 | 0.00008403922127 | -1.397911941 | Tubulin binding |
| CG6930 | l(3)neo38 | 0.000263 | -1.38 | NMJ development |
| CG6863 | tok | 0.004828954057 | -1.354579117 | Axonogenesis |
| CG3964 | TTLL4B | 0.00003165366938 | -1.329411974 | Tubulin binding |
| CG32435 | chb | 0.0000001207469809 | -1.301468318 | Tubulin binding |
| CG32435 | chb | 0.0000001207469809 | -1.301468318 | Axonogenesis |
| CG5539 | CG5539 | 0.0004105095793 | -1.292629299 | Tubulin binding |
| CG9308 | CG9308 | 0.006065418894 | -1.278143765 | Endocytosis |
| CG32796 | boi | 0.00001601031891 | -1.268457669 | Axonogenesis |
| CG9129 | Bfc | 0.0007192504581 | -1.246781338 | Endocytosis |
| CG17697 | fz | 0.00001032721276 | -1.211835294 | Axonogenesis |
| CG1765 | EcR | 0.002028601278 | -1.2064924 | Endocytosis |
| CG31641 | stai | 0.003627004825 | -1.20432594 | Tubulin binding |
| CG6433 | qua | 0.0001705390689 | -1.202132059 | Actin binding |
| CG31641 | stai | 0.00363 | -1.2 | NMJ development |
| CG30183 | CG30183 | 0.00003730926791 | -1.157736952 | Actin binding |
| CG10967 | Atg1 | 0.002051043358 | -1.154769282 | Axonogenesis |
| CG10967 | Atg1 | 0.00205 | -1.15 | NMJ development |
| CG16719 | CG16719 | 0.007284511382 | -1.134527758 | Tubulin binding |
| CG7210 | kel | 0.001107917663 | -1.131498886 | Actin binding |
| CG9211 | ihog | 0.001326885883 | -1.113938436 | Axonogenesis |
| CG393 | N | 0.008732747085 | -1.096549933 | Axonogenesis |
| CG14781 | mei-38 | 0.005217062424 | -1.088833247 | Tubulin binding |
| CG9554 | eya | 0.0008707210223 | -1.077573836 | Axonogenesis |
| CG15427 | tutl | 0.004493519437 | -1.075744853 | Axonogenesis |
| CG2988 | ems | 0.0007453389183 | -1.070996467 | Axonogenesis |
| CG32858 | sn | 0.007338953143 | -1.070761202 | Actin binding |
| CG7147 | kuz | 0.001569518294 | -1.051287446 | Axonogenesis |
| CG8590 | Klp3A | 0.0006115595912 | -1.048872231 | Tubulin binding |
| CG17348 | *drl* | 0.00003970382549 | -1.04017189 | Axonogenesis |
| CG3929 | *dx* | 0.001456360753 | -1.035176322 | Endocytosis |
| CG45781 | *DIP-lambda* | 0.00328 | -1.03 | NMJ development |
| CG6392 | *cmet* | 0.000005942658714 | -1.02624048 | Tubulin binding |
| CG13388 | *Akap200* | 0.001694057262 | -0.9863765658 | Actin binding |
| CG9218 | *sm* | 0.005443338125 | -0.9814585193 | Axonogenesis |
| CG8532 | *lqf* | 0.008242766109 | -0.9756690663 | Endocytosis |
| CG1655 | *sofe* | 0.001021447238 | -0.9748014683 | Tubulin binding |
| CG1193 | *kat-60L1* | 0.006369464376 | -0.9609529934 | Tubulin binding |
| CG6539 | *Gem3* | 0.00675 | -0.95 | NMJ development |
| CG11895 | *stan* | 0.007946024031 | -0.9433635179 | Axonogenesis |
| CG10718 | *neb* | 0.001755700466 | -0.9415672661 | Tubulin binding |
| CG16833 | *TTLL4A* | 0.001626598968 | -0.8884450639 | Tubulin binding |
| CG4166 | *not* | 0.006910267649 | -0.8567534233 | Axonogenesis |
| CG8376 | *ap* | 0.0003421629311 | -0.8234733892 | Axonogenesis |
| CG6875 | *asp* | 0.000530179956 | -0.787318857 | Tubulin binding |
| CG10923 | *Klp67A* | 0.00322270664 | -0.7633139696 | Tubulin binding |
| CG1848 | *LIMK1* | 0.00478 | -0.76 | NMJ development |
| CG1848 | *LIMK1* | 0.004777463437 | -0.7595434259 | Axonogenesis |
| CG6224 | *dbo* | 0.00166 | -0.757 | NMJ development |
| CG8705 | *pnut* | 0.009390097503 | -0.7434457115 | Actin binding |
| CG8705 | *pnut* | 0.009390097503 | -0.7434457115 | Tubulin binding |
| CG10346 | *Grip71* | 0.003951991021 | -0.7228332782 | Tubulin binding |
| CG6203 | *Fmr1* | 0.006056583189 | -0.6814038106 | Axonogenesis |
| CG6203 | *Fmr1* | 0.00606 | -0.681 | NMJ development |
| CG1210 | *Pdk1* | 0.00355 | -0.671 | NMJ development |
| CG1258 | *pav* | 0.009232237806 | -0.6506378617 | Tubulin binding |
| CG3637 | *Cortactin* | 0.007817504964 | -0.573856745 | Actin binding |
| CG3637 | *Cortactin* | 0.007817504964 | -0.573856745 | Endocytosis |
| CG1771 | *mew* | 0.008466387763 | 0.6454121039 | Axonogenesis |
| CG7438 | *Myo31DF* | 0.007036211569 | 0.6721348332 | Actin binding |
| CG7438 | *Myo31DF* | 0.007036211569 | 0.6721348332 | Endocytosis |
| CG8739 | *stmA* | 0.004034016019 | 0.6737250827 | Endocytosis |
| CG8948 | *Graf* | 0.002295630062 | 0.7002806062 | Endocytosis |
| CG9155 | *Myo61F* | 0.002261426157 | 0.7052714662 | Actin binding |
| CG9155 | *Myo61F* | 0.002261426157 | 0.7052714662 | Endocytosis |
| CG13531 | *CG13531* | 0.007907424895 | 0.7196454552 | Axonogenesis |
| CG3630 | *CG3630* | 0.001876354062 | 0.7701054415 | Actin binding |
| CG14994 | *Gad1* | 0.00426 | 0.775 | NMJ development |
| CG1081 | *Rheb* | 0.00491 | 0.806 | NMJ development |
| CG1081 | *Rheb* | 0.004909778225 | 0.8064999365 | Axonogenesis |
| CG5387 | *Cdk5alpha* | 0.001921196871 | 0.8077335929 | Axonogenesis |
| CG2715 | *Syx4* | 0.000991 | 0.83 | NMJ development |
| CG17947 | *alpha-Cat* | 0.003319871394 | 0.911620752 | Actin binding |
| CG18657 | *NetA* | 0.0003814093929 | 0.9310388515 | Axonogenesis |
| CG8732 | *AcsI* | 0.001800292961 | 0.9440080828 | Axonogenesis |
| CG6531 | *wgn* | 0.001986574594 | 0.9759866901 | Axonogenesis |
| CG31033 | *Atg16* | 0.00009457644496 | 0.9937423564 | Endocytosis |
| CG31352 | *Unc-115a* | 0.001748061909 | 0.9945388232 | Actin binding |
| CG31352 | *Unc-115a* | 0.001748061909 | 0.9945388232 | Axonogenesis |
| CG6502 | *E(z)* | 0.00004562014765 | 0.9952567007 | Axonogenesis |
| CG1967 | *p24-1* | 0.008424531593 | 0.9977489956 | Endocytosis |
| CG5020 | *CLIP-190* | 0.003337831289 | 1.010374543 | Actin binding |
| CG5020 | *CLIP-190* | 0.003337831289 | 1.010374543 | Tubulin binding |
| CG15861 | *CG15861* | 0.005319422315 | 1.01323568 | Endocytosis |
| CG9379 | *by* | 0.002241149583 | 1.026901521 | Actin binding |
| CG32683 | *CG32683* | 0.001693837258 | 1.041680775 | Endocytosis |
| CG16944 | *sesB* | 0.00333 | 1.06 | NMJ development |
| CG11064 | *apolpp* | 0.002957901393 | 1.110018532 | Tubulin binding |
| CG5907 | *Frq2* | 0.000243 | 1.15 | NMJ development |
| CG17352 | *Culd* | 0.001689733065 | 1.190343584 | Endocytosis |
| CG6584 | *SelR* | 0.0000001322433722 | 1.234846316 | Axonogenesis |
| CG4898 | *Tm1* | 0.007644941495 | 1.285698049 | Actin binding |
| CG6867 | *CG6867* | 0.004197195262 | 1.28616222 | Axonogenesis |
| CG32120 | *sens* | 0.002727046788 | 1.307841523 | Axonogenesis |
| CG7000 | *Snmp1* | 0.0009237727713 | 1.315413112 | Endocytosis |
| CG33115 | *NimB4* | 0.0007246135132 | 1.352093452 | Endocytosis |
| CG12002 | *Pxn* | 0.008829805506 | 1.378844475 | Endocytosis |
| CG3359 | *mfas* | 0.0000003824369147 | 1.481581912 | Axonogenesis |
| CG6622 | *Pkc53E* | 0.001586659527 | 1.5774131 | Endocytosis |
| CG4099 | *Sr-CI* | 0.0002991020021 | 1.577820585 | Endocytosis |
| CG11988 | *neur* | 0.002293308583 | 1.594630816 | Endocytosis |
| CG3588 | *CG3588* | 0.001451128421 | 1.648355166 | Actin binding |
| CG16857 | *bdI* | 0.003426133818 | 1.652047971 | Axonogenesis |
| CG32227 | *gogo* | 0.00002063828439 | 1.665442041 | Axonogenesis |
| CG9904 | *Seipin* | 0.0006685905849 | 1.673124514 | Axonogenesis |
| CG31839 | *NimB2* | 0.0007733369589 | 1.681345137 | Endocytosis |
| CG31019 | *CG31019* | 0.0001774858868 | 1.812426554 | Tubulin binding |
| CG11584 | *CG11584* | 0.004249530844 | 1.86393945 | Tubulin binding |
| CG43772 | *mwh* | 0.00003368903968 | 1.975531773 | Actin binding |
| CG45057 | *ringer* | 0.00000002385061229 | 2.006014928 | Tubulin binding |
| CG5637 | *nanos* | 0.00119 | 2.08 | NMJ development |
| CG8907 | *CG8907* | 0.0004386506421 | 2.089268428 | Actin binding |
| CG45110 | *tau* | 0 | 2.325277331 | Tubulin binding |
| CG31783 | *ninaD* | 0.0002408478531 | 2.855506204 | Endocytosis |
| CG7526 | *frac* | 0.00000004695231425 | 3.342239433 | Axonogenesis |
| CG42399 | *CG42399* | 0 | 4.576449346 | Tubulin binding |

